# Development, Structure, and Mechanism of Synthetic Antibodies that Target Claudin and *Clostridium perfringens* Enterotoxin Complexes

**DOI:** 10.1101/2022.01.01.474715

**Authors:** Benjamin J. Orlando, Pawel K. Dominik, Sourav Roy, Chinemerem Ogbu, Satchal K. Erramilli, Anthony A. Kossiakoff, Alex J. Vecchio

## Abstract

Strains of the Gram-positive bacterium *Clostridium perfringens* produce a two-domain enterotoxin (CpE) that afflict millions of humans and domesticated animals annually by causing prevalent gastrointestinal illnesses. CpE’s C-terminal domain (cCpE) binds cell surface receptors then its N-terminal domain restructures to form a membrane-penetrating *β*-barrel pore, which is toxic to epithelial cells of the gut. The claudin family of membrane proteins are the receptors for CpE, and also control the architecture and function of cell/cell contacts called tight junctions that create barriers to intercellular transport of solutes. CpE binding disables claudin and tight junction assembly and induces cytotoxicity via *β*-pore formation, disrupting gut homeostasis. Here, we aimed to develop probes of claudin/CpE assembly using a phage display library encoding synthetic antigen-binding fragments (sFabs) and discovered two that bound complexes between human claudin-4 and cCpE. We established each sFab’s unique modes of molecular recognition, their binding affinities and kinetics, and determined structures for each sFab bound to ~35 kDa claudin-4/cCpE in three-protein comprised complexes using cryogenic electron microscopy (cryoEM). The structures reveal a recognition epitope common to both sFabs but also that each sFab distinctly conforms to bind their antigen, which explain their unique binding equilibria. Mutagenesis of antigen/sFab interfaces observed therein result in further binding changes. Together, these findings validate the structures and uncover the mechanism of targeting claudin-4/cCpE complexes by these sFabs. Based on these structural insights we generate a model for CpE’s cytotoxic claudin-bound *β*-pore that predicted that these two sFabs would not prevent CpE cytotoxicity, which we verify *in vivo* with a cell-based assay. This work demonstrates the development and targeting mechanisms of sFabs against claudin/cCpE that enable rapid structural elucidation of these small membrane protein complexes using a cryoEM workflow. It further provides a structure-based framework and therapeutic strategies for utilizing these sFabs as molecular templates to target claudin/CpE assemblies, obstruct CpE cytotoxicity, and treat CpE-linked gastrointestinal diseases that cause substantial economic and quality of life losses throughout the world.

## INTRODUCTION

Tight junctions (TJs) are molecular gatekeepers that regulate transport of small molecules through the paracellular spaces between adjoining cells in endothelia and epithelia. To accomplish this function, TJs possess integral membrane proteins that self-assemble to simultaneously span both intracellular and paracellular spaces (1, 2). Of the numerous membrane proteins at TJs, the 27-member family of claudins comprise the major structural and function backbone of TJs, making them attractive targets to modulate TJ barriers therapeutically (3). Evolution has successfully accomplished this feat. Type F strains of the pathogenic Gram-positive bacterium *Clostridium perfringens* produce an enterotoxin (CpE) that binds claudins to dissociate TJs during cytotoxicity in the gut (4–6). In domesticated animals, CpE causes necrotic enteritis, colitis, and diarrhea (7, 8). In humans CpE causes enterotoxemia; is the 3^rd^ most prevalent foodborne illness in the United States causing an estimated ~$400 million annual economic burden; and is the source of a further 4+ million food poisoning cases worldwide—some resulting in death (7, 9–12). Unlike other human diseases caused by *Clostridium perfringens* toxins, CpE-associated ailments are not directly preventable nor treatable, and no vaccine exists against this food poisoning type (13). Because CpE is heat-resistant and produced by spore-formed *Clostridium perfringens*, cooking does little to reduce its pathogenicity (14). Therefore, sub-molecular details into CpEs mechanisms of claudin binding and dissociating TJ barriers are essential to elucidate CpE cytotoxicity and to develop therapeutic strategies for CpE-based diseases.

The C-terminal domain of CpE (cCpE) selectively targets claudins in the gut by recognizing a motif unique to these receptors, then binds them with low nanomolar affinities (15, 16). This claudin-bound CpE, *i.e*. “small complex”, then oligomerizes and its N-terminal domain structurally rearranges to form a membrane-penetrating and cytotoxic β-barrel pore (17, 18). The process of β-pore formation disables claudin/claudin interactions vital to TJ assembly and ultimately dissociates TJs causing paracellular leakage prior to CpE-induced cell death. Crystal structures of cCpE bound to receptor claudins have shed light on their interprotein interactions and have helped to inform structure-guided design of modified cCpEs used to detect or destroy cancer cells or to modulate the blood-brain barrier for drug delivery (19–27). Yet, a complete structural and mechanistic understanding of CpE dissociation of TJs and the process of cytotoxic β-pore formation remains elusive.

We intended to elucidate how CpE binds claudins and dissociates TJs by determining X-ray crystal structures of enterotoxins CpE or cCpE in complex with claudins but found crystallization to be a bottleneck. For most claudins, crystals did not form at all, while for those that formed crystals it required screening and optimizing hundreds over ~one year to determine structures resolved to 3-4 Å (22, 23). Using a phage display library encoding synthetic antigen-binding fragments (sFabs) we sought to discover molecules that target and bind complexes between enterotoxins and human claudin-4 (claudin-4). Our goal was to use sFabs to chaperone crystallization, improve initial diffraction, and increase structural throughput of this and other claudin/enterotoxin complexes (28, 29). Through this approach, we surmised, additional sFabs could be discovered that obstruct complex formation altogether and be useful in therapeutic development. During this process three sFabs were discovered, which we termed <u>C</u>pE <u>O</u>bstructing <u>P</u>roteins (COPs). Preliminary characterization of COPs revealed that COP-2 and COP-3 had properties amenable for structure determination of the claudin-4/cCpE complex. Ultimately, however, we determined a structure of this complex using a traditional crystallography workflow (23). But because sFabs have recently been shown effective for determining structures of small membrane proteins and complexes by cryogenic electron microscopy (cryoEM), we used COP-2, COP-3, and cryoEM to progress a novel workflow for higher throughput elucidation of claudin/enterotoxin structures (30–32).

Here, we qualitatively and quantitatively characterize COP-2 and COP-3 binding to claudin-4, cCpE, and CpE individually, and to claudin-4/enterotoxin complexes using biochemical and biophysical techniques. We also use cryoEM at 200 keV to determine 4-7 Å structures of each 50 kDa COP bound to ~35 kDa claudin-4/cCpE complexes in ~one month; and employ these structures to create models of the cytotoxic CpE β-pore. Our findings reveal the structural basis and COP-specific mechanisms of COP targeting of claudin-4/cCpE complexes. This research independently validates claudin-4/cCpE structures determined by X-ray crystallography and provides a structural framework for preventing CpE-mediated cytotoxicity by obstructing toxin/receptor binding. Moreover, it advances development of technologies and establishes a general approach for determining structures of other claudin/enterotoxin complexes at moderate resolutions rapidly using practical cryoEM instrumentation that can readily be expanded to higher resolutions with 300 keV microscopes. Further, it demonstrates the antigenicity of CpE and cCpE enterotoxins and their claudin-bound complexes, which through intensified sFab development could generate novel sFabs useful for modulating TJ barriers or as therapeutics for preventing or treating CpE-based illnesses in humans and domesticated animals.

## RESULTS

### Development of COPs

claudin-4 solubilized in n-dodecyl-β-D-maltopyranoside (DDM) and bound to cCpE was used as input for phage display selection using a large and diverse library of sFabs based on a humanized Fab scaffold (see **Methods**). The sFab library has varied sequences that are biased for serine and tyrosine in the complimentarity-determining regions (CDRs) of their light (L) and heavy (H) chains within variable domains (33, 34). After several rounds of selection to increase stringency, two sFabs, termed COP-2 and COP-3, were further developed and validated by ELISA to bind to claudin-4/cCpE. Both COPs were sequenced, recombinantly expressed in *E. coli*, and purified using affinity chromatography in order to isolate these sFabs and characterize their binding further. Sequence alignments of COP-2 and COP-3 reveal their unique primary sequences (***SI Appendix***, **Fig. S1**). The alignments show that COP-2 and COP-3 share 98.2 and 93.3% sequence identity in their L and H chains, respectively. The COP sequences diverge the greatest in CDR-L3, CDR-H1, and CDR-H3. These residue divergences may direct COP-specific recognition of claudin-4/cCpE complexes.

### Biochemical Characterization of COPs

To obtain more detailed insights into COP recognition we determined which molecule COPs bind and if they use unique or common epitopes. After expressing and purifying claudin-4, cCpE, CpE, and COPs, increases in molecular masses as assessed by decreases in peak retention times with size-exclusion chromatography (SEC) was used to qualitatively characterize COP-2 and COP-3 binding modes (see **Methods**). We observed, after incubating COP-2 and COP-3 with claudin-4 and cCpE alone, that COPs did not bind claudin-4 but did form larger complexes with cCpE (***SI Appendix*, Fig. S2A** and **S2B**). Incubating claudin-4, cCpE, and COPs together showed that cCpE binds claudin-4 and that both COPs bound claudin-4/cCpE complexes (***SI Appendix*, Fig. S2C**). No mass increases were observed for COPs incubated with CpE (***SI Appendix*, Fig. S2D**). Lastly, incubating claudin-4, CpE, and COPs together showed that while CpE binds claudin-4 and forms a “small complex”, the COPs do not bind “small complexes” (***SI Appendix*, Fig. S2E**). To verify complex formation, peaks from SEC were pooled and subjected to SDS-PAGE, which showed the presence of individual proteins from associated complexes (***SI Appendix*, Fig. S2F**). To determine if COP-2 and COP-3 share a binding epitope, we incubated both together with cCpE. If COPs bound distinct epitopes a molecular mass shift greater than the individual cCpE/COP complex would result. SEC revealed no additive mass shift with both COPs present (***SI Appendix*, Fig. S2B**). These biochemical results suggest the specific molecular recognition of COPs.

### Biophysical Characterization of COPs

After qualitatively establishing COP binding we quantitated the affinities and kinetics of COP interactions with claudin-4 and enterotoxins. We determined the second-order association rate constant (k_on_), first-order dissociation rate constant (k_off_), and equilibrium dissociation constant (K_D_) of these interactions using bio-layer interferometry (BLI) (**Table 1**). BLI measurements were made using pre-formed claudin-4/enterotoxin complexes or cCpE alone in DDM, replicating the conditions of the phage display selections. For COP-2 and COP-3 binding to claudin-4/cCpE complexes we measured K_Ds_ of 52.3 and 98.4 nM, respectively. Comparing the binding rates revealed that COP-3 had 1.9- and 3.5-fold faster k_on_ and k_off_ rates compared to COP-2. The K_Ds_ of COP-2 and COP-3 to cCpE were 67.6 and 137.6 nM, respectively. Like COP-3 binding to claudin-4/cCpE complexes, the k_on_ and k_off_ rates were 1.4- and 2.1-fold faster when compared to COP-2. Finally, we measured K_Ds_ of 7.8 and 9.1 μM for COP-2 and COP-3 binding to claudin-4/CpE “small complexes”, respectively. The K_D_ values represent 149.1- and 92.0-fold decreases in COP-2 and COP-3 affinity for CpE compared to cCpE. Kinetic differences in k_on_ and k_off_ of COP binding are visible in the BLI sensorgrams (**Fig. 1B, 2B**, and ***SI Appendix*, Fig. S3**). These results agree with biochemical assessment and reveal biophysical parameters unique to each COP that may influence recognition and binding to claudin-4/cCpE complexes.

**Figure 1.**
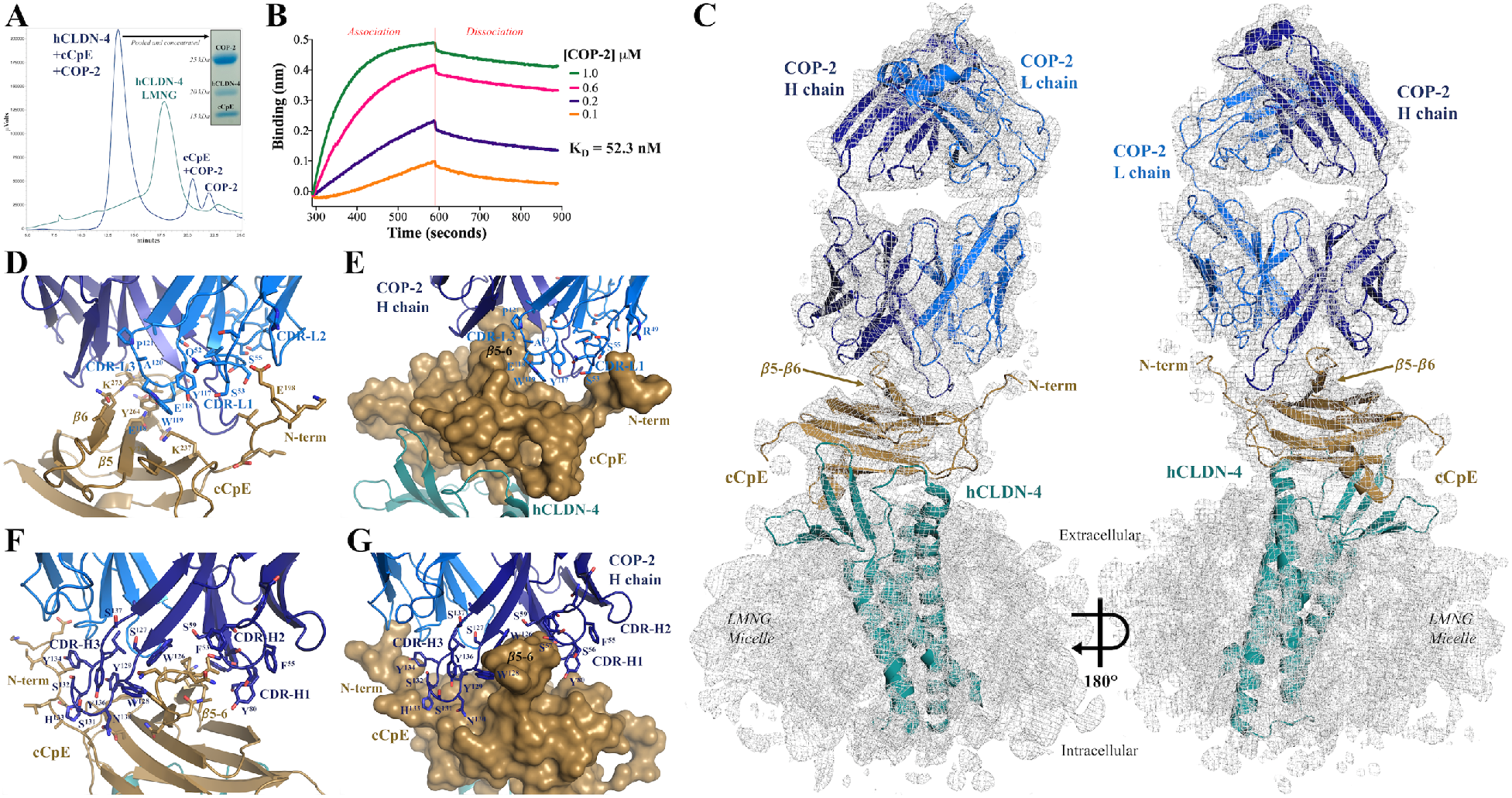
Structure and Function of COP-2 Binding to Claudin-4/cCpE Complexes. (**A**) SEC purification of human claudin-4 (hCLDN-4)/cCpE/COP-2 complexes in LMNG detergent. (**B**) Binding of COP-2 to claudin-4/cCpE using bio-layer interferometry. (**C**) CryoEM structure of COP-2 (blue) bound to cCpE (copper) and hCLDN-4 (teal). Proteins are shown as cartoons within the cryoEM map (grey), which is contoured to 4.0 σ. (**D-G**) Potential interactions between COP-2 (blue) and cCpE (copper) for: (**D** and **E**) Chain L (light blue); and (**F** and **G**) Chain H (dark blue). COP-2 and cCpE are both represented as cartoons (**D** and **F**) or, COP-2 as a cartoon and cCpE as a surface (**E** and **G**).

**Figure 2.**
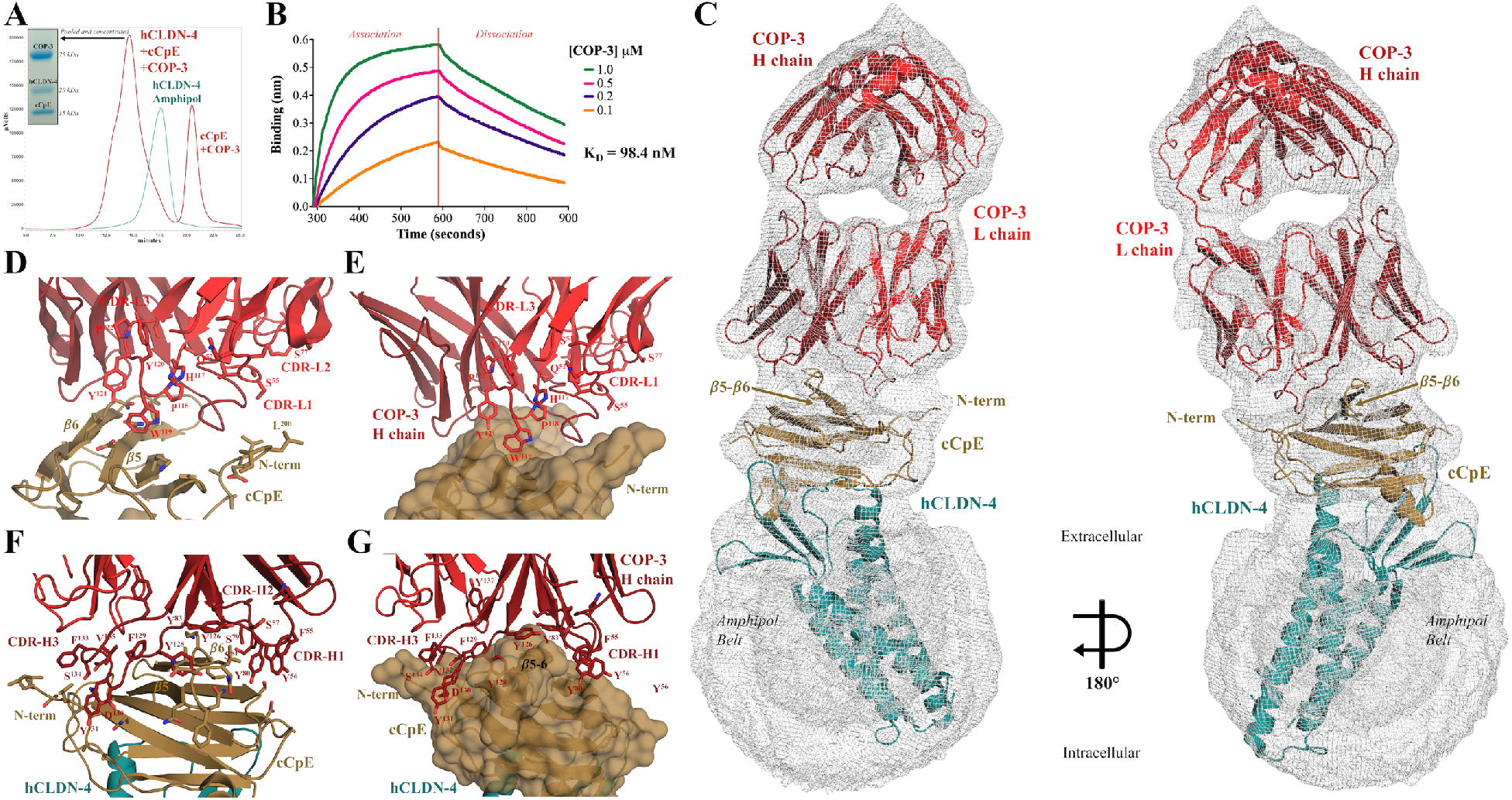
Structure and Function of COP-3 Binding to Claudin-4/cCpE Complexes. (**A**) SEC purification of human claudin-4 (hCLDN-4)/cCpE/COP-3 complexes in amphipol A8-35. (**B**) Binding of COP-3 to claudin-4/cCpE using bio-layer interferometry. (**C**) CryoEM structure of COP-3 (red) bound to cCpE (copper) and hCLDN-4 (teal). Proteins are shown as cartoons within the cryoEM map (grey), which is contoured to 5.0 σ. (**D-G**) Potential protein/protein interactions between COP-3 (red) and cCpE (copper) for: (**D** and **E**) Chain L (red); and (**F** and **G**) Chain H (maroon). COP-3 and cCpE are both represented as cartoons (**D** and **F**) or, COP-3 as a cartoon and cCpE as a surface (**E** and **G**).

**Table 1:** Affinities and Kinetics of COP Binding to Claudin-4/cCpE and Enterotoxins.

| **COP** | **Complex/Protein** | **$k_{on}$ (1/Ms)** | **$k_{off}$ (1/s)** | **$K_D$ (nM)** |
| --- | --- | --- | --- | --- |
| **COP-2** | claudin-4/cCpE | $1.5 \times 10^4$ | $7.8 \times 10^{-4}$ | **52.3** |
| | | $\pm 0.1 \times 10^4$ | $\pm 0.6 \times 10^{-4}$ | $\pm 3.0$ |
| | cCpE | $7.2 \times 10^3$ | $4.8 \times 10^{-4}$ | **67.6** |
| | | $\pm 0.6 \times 10^3$ | $\pm 0.4 \times 10^{-4}$ | $\pm 11.7$ |
| | claudin-4/CpE | $1.5 \times 10^2$ | $1.2 \times 10^{-3}$ | **7,800.0** |
| | | $\pm 0.6 \times 10^2$ | $\pm 0.6 \times 10^{-3}$ | $\pm 750.0$ |
| **COP-3** | claudin-4/cCpE | $2.9 \times 10^4$ | $2.7 \times 10^{-3}$ | **98.4** |
| | | $\pm 0.7 \times 10^4$ | $\pm 0.5 \times 10^{-3}$ | $\pm 11.0$ |
| | cCpE | $1.0 \times 10^4$ | $1.0 \times 10^{-3}$ | **137.6** |
| | | $\pm 0.1 \times 10^4$ | $\pm 0.1 \times 10^{-3}$ | $\pm 6.2$ |
| | claudin-4/CpE | $1.2 \times 10^3$ | $10.8 \times 10^{-2}$ | **9,050.0** |
| | | $\pm 0.2 \times 10^3$ | $\pm 0.7 \times 10^{-2}$ | $\pm 1,300.0$ |

### CryoEM Structure of claudin-4/cCpE/COP-2 Complex

Having confirmed that COP-2 binds claudin-4/cCpE complexes using SEC (**Fig. 1A**) and BLI (**Fig. 1B**), we next delineated the structural basis and molecular mechanism of COP-2 targeting by determining its structure by cryoEM. CryoEM screening of claudin-4/cCpE/COP-2 complexes that were SEC purified, pooled, and concentrated to 8.0 mg/mL in various detergents showed that those solubilized in 2,2-didecylpropane-1,3-bis-β-D-maltopyranoside (LMNG) were amenable to structure determination. This was confirmed by subsequent 2D classifications and *ab initio* 3D reconstructions, which revealed a stacked and linear arrangement of the three proteins, a canonical two-lobed sFab, and flexibility in COP-2’s constant domain (***SI Appendix***, **Fig. S4**). Due to this flexibility, we used a data processing strategy that masked claudin-4, cCpE, and the variable domains of COP-2 to resolve and focus understanding on the interactions directing COP-2 recognition. Using this strategy, we generated a final processed cryoEM map that was resolved to 6.9 Å (***SI Appendix*, Table S1**). The map resolution was sufficient to reveal the claudin-4/cCpE/COP-2 complex, secondary structural elements including claudin-4’s four transmembrane helices (TM), and some bulky side chains (***SI Appendix*, Fig. S5A**). This cryoEM map was used to build, refine, and determine a structure for the claudin-4/cCpE/COP-2 complex. ***SI Appendix*, Table S1** shows data processing, refinement, and model-to-map fit statistics, with further details given in **Methods**.

The cryoEM map resolution was insufficient to place side chains confidently, so we cannot verify interactions between claudin-4 and cCpE that define the “cCpE-binding motif” (23). Overall, however, the claudin-4/cCpE portion of this complex from cryoEM superimposes well onto the crystal structure (***SI Appendix*, Fig. S6A**). We measured root mean square deviations (RMSDs) in Cα positions of 2.0 and 1.4 Å between the cores of claudin-4 and cCpE, and 1.7 Å in overall secondary structures between the cryoEM and crystal structures of the claudin-4/cCpE complex, indicating no major conformational changes occur upon COP-2 binding. Generally, the claudin-4 extracellular segments (ECS) are in similar conformations and thus may interact with cCpE similarly in the cryoEM and crystal structures (***SI Appendix*, Fig. S6A**). Density corresponding to a loop within ECS2, which contains the NPLVA^153^ motif shows that the motif accesses a groove on the surface of cCpE. This interaction is known to impart high-affinity cCpE binding to claudins and COP-2 appears to not significantly alter its structure (23).

The cryoEM structure of the claudin-4/cCpE/COP-2 complex reveals the basis of COP-2 binding. The canonical binding of cCpE to claudin-4 exposes cCpE’s top half to COP-2 binding by providing an antigenic surface (**Fig. 1C**). COP-2 binding to cCpE alters the conformations of cCpE’s N-terminus, *β*-strands *β*5 and *β*6, and the loop connecting them, when compared to the crystal structure (***SI Appendix*, Fig. S6A**). COP-2’s L chain sits atop a depression on the exterior of cCpE formed between the N-terminus and strands *β*5 and *β*6 (**Fig. 1C**). Chain L’s CDR-L1, -L2, and -L3 conform to the surface of cCpE and potential side chains involved in these interactions can be visualized although not placed confidently (**Fig. 1D** and **1E**). COP-2’s chain H access the same surface depression as chain L but also flanks the opposite side of strands *β*5 and *β*6—CDR-H3 shares an epitope with chain L while CDR-H1 and -H2 reside on the other (**Fig. 1C** and **1F**). CDR-H3 splays outward to deeply penetrate its surface groove, conforming to the cCpE surface (**Fig. 1G**). Based on the interactions projected by the structure, we hypothesized that residues comprising Lys197 to Leu202 and Asn267 to Gln276 in cCpE could influence COP-2 binding.

### CryoEM Structure of claudin-4/cCpE/COP-3 Complex

After verifying that COP-3 binds claudin-4/cCpE complexes using SEC (**Fig. 2A**) and BLI (**Fig. 2B**), we next determined a structure for COP-3 in complex with claudin-4/cCpE by cryoEM to contrast its molecular mechanism of targeting with COP-2. claudin-4/cCpE/COP-3 complexes in various membrane mimetics were SEC purified, pooled, and concentrated to 6.0 mg/mL for cryoEM (see **Methods**). Those complexes solubilized in amphipol were superior to LMNG and DDM for cryoEM based on 2D class averages. The 2D and 3D classifications of the COP-3 complex showed a single complex with each protein stacked in a linear arrangement and had features corresponding to secondary structural elements and both lobes of COP-3, indicating that maps of adequate resolution were obtained (***SI Appendix***, **Fig. S7**). The 3D reconstructions showed that the constant domains of COP-3 are less dynamic than those of COP-2, but also that claudin-4’s TM region was less well resolved in amphipol. We used a data processing strategy that masked COP-3 to best resolve the interactions directing its recognition of claudin-4/cCpE, which generated a final processed map *focused*) that was resolved to 3.8 Å and showed structural features including density for bulky side chains (***SI Appendix*, Table S1**). However, this map lacked definition in the cCpE and claudin-4 regions, and optimization of the model-to-map fit proved difficult. Therefore, data was processed with no mask, which produced a final map (*whole*) that was resolved to 5.0 Å. The *whole* map resolved the claudin-4/cCpE/COP-3 complex, including secondary structural elements like claudin-4’s TMs and α-helices and *β*-strands in cCpE, and the conformations of COP-3 CDRs (***SI Appendix*, Fig. S5B**). Initial model building and refinement employed both maps, but the final model was refined against the *whole* map, resulting in the cryoEM structure for the claudin-4/cCpE/COP-3 complex. ***SI Appendix*, Table S1** shows data processing, refinement, and model-to-map fit statistics, with further details given in **Methods**.

The cryoEM structure of the claudin-4/cCpE/COP-3 complex reveals COP-3’s mode of binding. Compared to the claudin-4/cCpE crystal structure, the equivalent portion from the COP-3 cryoEM structure superimposes well and indicates that COP-3 does not induce large conformational changes or affect normal claudin-4/cCpE interactions (***SI Appendix*, Fig. S6B**). We measured RMSDs in Cα positions of 2.2 and 1.2 Å between the cores of claudin-4 and cCpE, and 2.1 Å between overall secondary structures in the claudin-4/cCpE complex when comparing cryoEM and X-ray structures. Like COP-2, COP-3 accesses cCpE’s surface opposite to where claudin-4 binds due their canonical interactions (**Fig. 2C**). COP-3 binding to cCpE alters cCpE’s N-terminus, *β*5 and *β*6, and the loop connecting them, when compared to the crystal structure (***SI Appendix*, Fig. S6B**). The L chain of COP-3 binds between cCpE’s N-terminus and *β*5 and *β*6 using CDR-L1 and CDR-L3 (**Fig. 2D**). The CDR-L3 loop conforms to the surface of this region, potentially using aromatic side chains to drive shape complementarity (**Fig. 2E**). COP-3’s H chain flanks both sides of *β*5 and *β*6 of cCpE with CDR-H3 sharing an epitope with chain L while CDR-H1 and CDR-H2 occupy the other side (**Fig. 2C** and **2F**). COP-3’s CDR-H3 conforms to and deeply accesses a surface groove between this region and the N-terminus (**Fig. 2G**). Based on the interactions approximated by the structure, we hypothesized that residues comprising Glu198 to Leu202 and Asn269 to Gln276 in cCpE may guide COP-3 binding.

### Comparison of COP Structures

Because structure resolution was limiting we used computation-based structure and sequence analyses to estimate the COP-specific residues used for cCpE recognition. For this, we input our structures into PDBePISA, an online tool for determining the structural and chemical properties of macromolecular interfaces, and then compared areas determined by PDBePISA to preside over cCpE/COP binding with COP primary sequences (35) (***SI Appendix*, Fig. S1**). For COP-2 we found that CDR-L3 residues Tyr117 to Ala120, CDR-H1 residue Ser56, and CDR-H3 residues Tyr125 to Ser137 were unique to COP-2. For COP-3 we found that CDR-L3 residues His117 to Tyr121, CDR-H1 residue Tyr56, and CDR-H3 residues Gly125 to Tyr137 were unique to COP-3. When compared to a generic sFab, COPs contain many aromatic side chains and are enriched in serine and tyrosine residues in their CDRs (***SI Appendix*, Fig. S1**). This analysis exposes potential COP amino acid determinants for recognition and binding of claudin-4/cCpE complexes.

To contrast COP binding modes and to explain their varied biophysical binding equilibria, we overlaid our two cryoEM structures (***SI Appendix*, Fig. S8**). The overlays revealed that: 1) the cCpE poses when bound to claudin-4 are similar but structural perturbations exist in cCpE due to COP-induced changes; 2) the claudin-4/cCpE portions exhibit minor structural differences between complexes but the claudin-4 TMs are oriented differently in membrane mimetics; and 3) the COPs have similar secondary structural elements, but their tertiary structures and CDRs vary in conformations (***SI Appendix*, Fig. S8A**). Focusing on regions with the largest observable differences, we found that the L and H chains of each COP conform to cCpE uniquely. For COP-2 chain L, CDR-L1 and CDR-L3 reside within to the surface groove formed between the N-terminus and *β*5-*β*6 of cCpE while CDR-L2 lies external (***SI Appendix*, Fig. S8B**). For COP-3 chain L, the surface groove is depressed due to less N-terminal length, so while CDR-L1 and CDR-L3 reside in the same groove, CDR-L1 appears to interact less with cCpE. The conformation of CDR-H3 may force this CDR-L1 change in COP-3. Using PDBePISA we calculated interface surface areas between the L chain of COPs and cCpE and found that COP-2’s area was 36.4% larger. In both COP’s chain H, the CDR-H1 and CDR-H2 bind on one side of cCpE’s *β*5-*β*6 element while CDR-H3 flanks the other (***SI Appendix*, Fig. S8C**). CDR-H1 and CDR-H2 have similar conformations and sequence alignments show that residue conservation is high here (***SI Appendix*, Fig. S1**). But CDR-H3 conformations appear to vary and sequences also show the greatest diversity exists here. Generally, however, the conformations of all three CDRs overlay well between COPs, which is reflected by cCpE/H chain interface surface area differences of only 4.5%. For both COPs, ~80% of the total cCpE/COP interface area resides on chain H. Overall, COP-2 appears to employ chain L to a higher degree than COP-3, while both COPs use chain H similarly and more dominantly to chain L to bind cCpE. Moreover, the surface structure of cCpE appears to be changed more by COP-2 than COP-3 as a result of binding, indicating that COP-specific interactions, driven by sequence diversity, may uniquely mold cCpE’s surface (***SI Appendix*, Fig. S8B** and **S8C**). We next determined whether the structural observations detailed here were true *in vitro* and if they could explain the measured differences in binding equilibria between COP-2 and COP-3.

### Quantification of COP Binding to Mutant cCpEs

To test our structures and pinpoint the amino acid determinants of COP binding we mutated regions of cCpE where we observed potential interactions with COPs and quantified binding. **Table 2** shows sequences for cCpE^mutants^ that were generated. Pre-formed claudin-4/cCpE^mutant^ complexes were used to mimic the sFab selection experiment. First, we found that no cCpE^mutant^ affected binding to claudin-4, which binds with a K_D_ of 2.5 nM (***SI Appendix*, Fig. S3E**) (23, 36). Next, we tested COP binding to claudin-4/cCpE^mutant^ complexes using BLI (**Table 3**). Binding measurements showed that mutant cCpE^1^, which has a shortened N-terminus, decreased affinity for COP2 and COP-3 primarily by increasing k_off_ rates. Mutant cCpE^2^ increased COP-2 and COP-3 affinities through increased k_on_ rates—this mutation lies at the end of the N-terminus at the start of globular cCpE. Mutant cCpE^2^ with an added Leu202Ala mutation, cCpE^2L^, showed binding to both COP-2 and COP-3 at near cCpE^wild type^ affinities, indicating a loss of affinity compared to mutant cCpE^2^ due to the shorter alanine side chain. Kinetics reveal that COPs bind claudin-4/cCpE^2L^ with slower k_on_ and faster k_off_ rates compared to claudin-4/cCpE^2^. The mutant cCpE^3^ in the loop connecting *β*5 to *β*6 deceased affinity for COP-2 by 1.2-fold—but for COP-3 it deceased affinity 12.4-fold—this greater affinity loss for COP-3 is driven by a 14.4-fold faster k_off_ rate. Both COPs exhibited substantial losses in affinity to mutant cCpE^4^, which alters residues in the *β*5 to *β*6 connecting loop and *β*6 to alanine. COP-2 affinity decreased 4,835.6-fold while COP-3 decreased 135.2-fold. Unlike other cCpE mutants, cCpE^4^ affected both k_on_ and k_off_ rates, indicating that this mutant radically perturbed normal COP binding. To put these results in a structural context, we made models of mutant cCpEs based on our cryoEM structures in order to compare them to cCpE^wildtype^ (***SI Appendix*, Fig. S9**). These models provide structural bases for COP binding to mutant cCpEs that explain our biophysical measurements. Overall, each set of cCpE mutants, chosen based on their potential for side chain interactions with COPs, altered COP binding in different ways, providing *in vitro* validation of our structures.

**Table 2:**
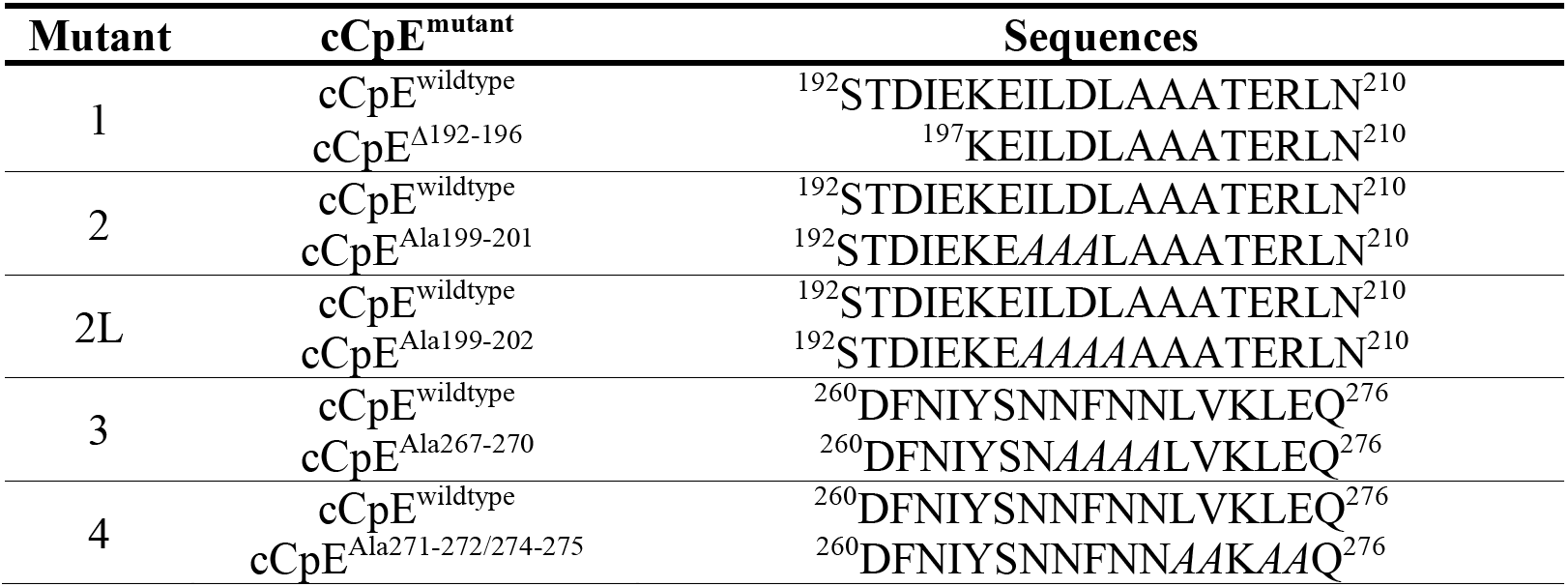
Mutant cCpE Sequences.

**Table 3:** Affinities and Kinetics of COP Binding to Claudin-4/cCpE^mutant^ Complexes.

| **COP** | **claudin-4/cCpE<sup>mutant</sup>** | **k<sub>on</sub> (1/Ms)** | **k<sub>off</sub> (1/s)** | **K<sub>D</sub> (nM)** |
| --- | --- | --- | --- | --- |
| **COP-2** | cCpE<sup>wildtype</sup> | 1.5x10<sup>4</sup><br>± 0.1x10<sup>4</sup> | 7.8x10<sup>-4</sup><br>± 0.6x10<sup>-4</sup> | **52.3**<br>± 3.0 |
|  | cCpE<sup>1</sup> | 7.7x10<sup>3</sup><br>± 0.5x10<sup>3</sup> | 1.2x10<sup>-3</sup><br>± 0.1x10<sup>-3</sup> | **161.3**<br>± 0.6 |
|  | cCpE<sup>2</sup> | 3.9x10<sup>4</sup><br>± 0.5x10<sup>4</sup> | 3.3x10<sup>-4</sup><br>± 0.4x10<sup>-4</sup> | **8.6**<br>± 0.1 |
|  | cCpE<sup>2L</sup> | 1.7x10<sup>4</sup><br>± 0.1x10<sup>4</sup> | 1.0x10<sup>-3</sup><br>± 0.1x10<sup>-3</sup> | **57.2**<br>± 3.7 |
|  | cCpE<sup>3</sup> | 1.3x10<sup>4</sup><br>± 0.1x10<sup>4</sup> | 9.4x10<sup>-4</sup><br>± 0.1x10<sup>-4</sup> | **64.5**<br>± 4.3 |
|  | cCpE<sup>4</sup> | 5.3x10<sup>1</sup><br>± 1.2x10<sup>1</sup> | 1.3x10<sup>-2</sup><br>± 0.1x10<sup>-2</sup> | **252,900.0**<br>± 40,700.0 |
| **COP-3** | cCpE<sup>wildtype</sup> | 2.9x10<sup>4</sup><br>± 0.7x10<sup>4</sup> | 2.7x10<sup>-3</sup><br>± 0.5x10<sup>-3</sup> | **98.4**<br>± 11.0 |
|  | cCpE<sup>1</sup> | 1.9x10<sup>4</sup><br>± 0.2x10<sup>4</sup> | 3.9x10<sup>-3</sup><br>± 0.6x10<sup>-3</sup> | **200.2**<br>± 5.5 |
|  | cCpE<sup>2</sup> | 11.0x10<sup>4</sup><br>± 2.2x10<sup>4</sup> | 3.0x10<sup>-3</sup><br>± 0.1x10<sup>-3</sup> | **24.6**<br>± 4.7 |
|  | cCpE<sup>2L</sup> | 6.3x10<sup>4</sup><br>± 0.2x10<sup>4</sup> | 5.3x10<sup>-3</sup><br>± 0.1x10<sup>-3</sup> | **84.4**<br>± 4.8 |
|  | cCpE<sup>3</sup> | 3.0x10<sup>4</sup><br>± 0.3x10<sup>4</sup> | 3.9x10<sup>-2</sup><br>± 1.1x10<sup>-2</sup> | **1224.6**<br>± 143.7 |
|  | cCpE<sup>4</sup> | 1.0x10<sup>3</sup><br>± 0.1x10<sup>3</sup> | 1.3x10<sup>-2</sup><br>± 0.1x10<sup>-2</sup> | **13,300.0**<br>± 10.0 |

### Model of Cytotoxic Claudin-bound CpE Pore

To predict COP function in the context of CpE-induced cytotoxicity, we created models for the process of COP binding to the claudin-4/CpE “small complex” and for a claudin-4-bound CpE β-barrel pore complex, using our cryoEM structures as guides (*see* **Methods**). We first modeled CpE binding to claudin-4 then COP recognition of the “small complex” (**Fig. 3A**). The model shows that if CpE is bound to claudin-4 in a “small complex” then its N-terminal domain would sterically shield COPs from access to their binding epitopes on cCpE. This finding explains results we obtained from biophysical measurements (**Table 1**). We next modelled the claudin-4-bound CpE β-pore complex (**Fig. 3B**). As there is no structure for the CpE β-pore, we used the cryoEM structure of lysenin, a β-pore toxin with homology to CpE to model it (37). Lysenin was chosen because attempts to model the complex using the aerolysin β-pore, another homologous protein, did not place claudin perpendicular to the membrane plane. We thus hypothesize that the CpE β-pore resembles lysenin more than aerolysin. This model reveals that when oligomeric CpE assembles into a β-pore complex that COP access to cCpE binding epitopes would be sterically obstructed even if CpE N-terminal domain rearrangements occur (**Fig. 3B** and **3C**). These structural models explain our *in vitro* binding data and provide a prediction for the effect of COPs on CpE-induced cytotoxicity *in vivo*.

**Figure 3.**
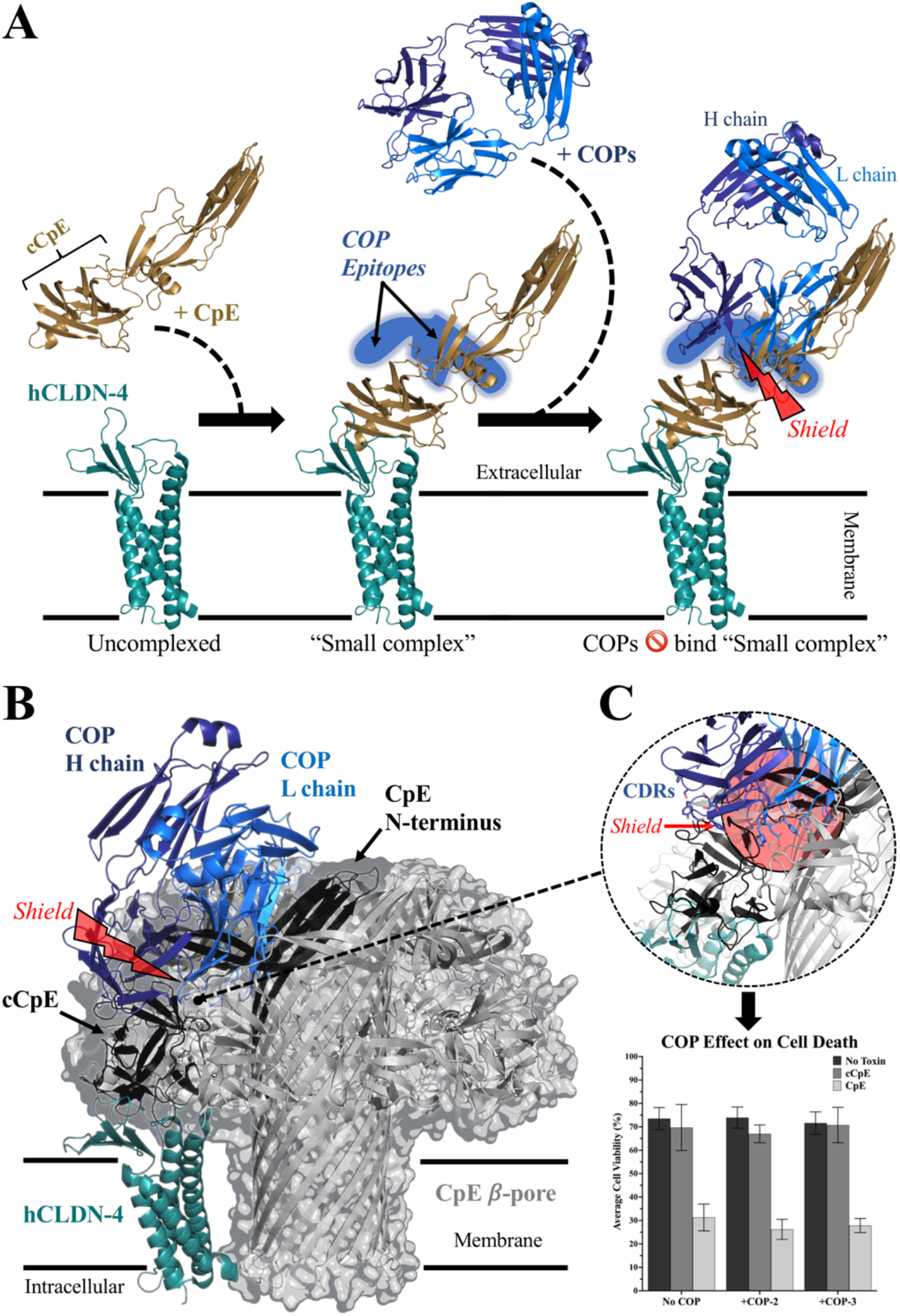
Models for COP Binding to Claudin-4/cCpE and the Claudin-4-bound CpE β-Pore. (**A**) CpE (copper) binding to the ECS of human claudin-4 (hCLDN-4, teal) forms the “small complex”. The N-terminus of CpE sterically shields the COP (blue) binding epitopes on cCpE, preventing high-affinity interactions. (**B**) Model for a CpE *β*-pore (grey) based on the structure of lysenin with one hCLDN-4 (teal) bound to the cCpE domain of a CpE monomer (black). Structural rearrangements of the CpE N-terminus forms the *β*-pore. Despite this rearrangement, CpE multimeric assembly and the N-terminus of CpE sterically obstruct full COP (blue) engagement with its binding epitopes. (**C**) Zoom-in on the sterically shielded region shows CDRs from the L and H chains cannot access their epitopes on cCpE. Associated bar graph shows the functional effect of COP addition to CpE-induced cytotoxicity of insect cells expressing claudin-4 on their surfaces. Graph plots the mean and standard deviation based on four readings. Proteins are represented as cartoons (**A** and **B**), or as a translucent surface encapsulated cartoon in the case of the CpE *β*-pore (**B**).

### Effect of COPs on CpE Cytotoxicity

Finally, we used a cell-based assay to validate our model of the claudin-4-bound CpE β-pore and to test whether COPs affect CpE-induced cytotoxicity (see **Methods**). Using *Spodoptera frugiperda* cells that lack endogenous claudins but form tight junction-like strands when expressing claudin-4 on their cell surfaces, we added COP-2 or COP-3, followed immediately by cCpE or CpE, using previously described methods (22, 23, 38, 39). Control wells consisted of no addition of COPs or enterotoxins. We then measured cell viability by quantifying the amount of cell death instigated by enterotoxins to determine whether COPs altered cytotoxicity (**Fig. 3C**). For cells expressing claudin-4 alone not treated with COPs we found that cell viability averaged 73.5%. For these cells, COP addition decreased average viability by 0.7%, indicating no COP-induced cytotoxicty. For cells expressing claudin-4 and treated with cCpE, which lacks the cytotoxic N-terminus, we found that cell viability averaged 69.7%. Again, for cCpE-treated cells COP addition decreased average viability only 0.8%. Finally, for cells expressing claudin-4 treated with CpE we found that cell viability averaged 31.3%, a decrease of 40.3% compared to untreated and cCpE-treated cells, indicating CpE-induced cytotoxicity. Addition of COPs to CpE-treated cells did not significantly change CpE-induced cell death, decreasing average cell viability by 4.3%. These results validate our *in silico* model of the claudin-4-bound CpE β-pore and establish the effect of COPs to CpE-induced cytotoxicity.

## DISCUSSION

Here we demonstrate the development of sFabs called <u>C</u>pE <u>O</u>bstructing <u>P</u>roteins, COP-2 and COP-3. We show that COPs bind well to cCpE but not to claudin-4 or CpE; that COPs bind cCpE better when bound to claudin-4 than when alone in solution; and that both COPs bind the same surface epitope on cCpE opposite to where claudin-4 binds (***SI Appendix*, Fig. S2**). We also show that COP binding to claudin-4/cCpE complexes yield similar affinities but different kinetics, with COP-2 associating and dissociating more slowly than COP-3 (**Table 1**). As COPs target and bind cCpE on the opposing surface to its claudin binding motif they are therefore capable of binding to any claudin/cCpE complex, making them a general yet strategic tool for enabling structures of claudin/cCpE complexes.

As proof of this concept, we determine cryoEM structures for COP-2 and COP-3 bound to claudin-4/cCpE complexes, which reveal the cCpE binding epitope and the potential interaction interfaces for the L and H chains of COPs (**Figs. 1** and **2**). Our structures and computational analyses show that COP-2 has an 11% larger cCpE/COP interface area, that COP-2 uses chain L CDR-L1 uniquely, and that its binding conforms to or molds cCpEs surface to a greater extent than COP-3, which explains its higher affinity and slower association and dissociation rates compared to COP-3. These analyses also reveal a common binding property to COPs where chain H interacts with both sides of a *β*5-*β*6 epitope in a mechanism similar to a caliper brake on a bicycle (**Figs. 1G** and **2G**). Holistically, the cryoEM structures resemble and thus provide independent validation of the claudin-4/cCpE complex crystal structure (***SI Appendix*, Fig. S6**).

To validate these moderate resolution cryoEM structures we mutate the observed COP binding epitopes on cCpE and quantify mutant effects to COP binding. For this we mutated side chains in sequential three to five residue zones, rather than making single point mutations (**Table 2**). We show that all mutations made and tested affect cCpE/COP binding affinity, kinetics, or both (**Table 3**). Some, like mutants cCpE^3^ and cCpE^4^, display COP-specific differences in their binding and thus pinpoint residues of cCpE that are uniquely recognized by either COP-2 or COP-3. The changes to COP binding by cCpE mutants *in vitro* validates our cryoEM structures by attesting to the accuracy of their modeled cCpE/COP interfaces. We further show that models of mutant cCpE/COP complexes guided by our structures elucidate the structural bases of mutant effects to COP binding (***SI Appendix*, Fig. S9**). In sum, these results provide the likely amino acid determinants and biophysical interaction mechanisms that direct COP-specific recognition of and binding to cCpE.

We further present models for the claudin-4/CpE “small complex” and claudin-4-bound CpE β-pore complex in order to understand our structures in a functional context and to predict COP efficacy for use as <u>C</u>pE <u>O</u>bstructing <u>P</u>roteins (**Fig. 3**). One model shows that when CpE is bound to claudin-4 in a “small complex” that the N-terminus of CpE sterically shields COP binding epitopes and prevents binding (**Fig. 3A**). This model is verified by our finding that COP-2 and COP-3 bound CpE with ~100-fold lower affinity than cCpE (**Table 1**). Using the homologous β-pore-forming toxin lysenin as a benchmark, we also show how CpE may oligomerize to form its plasma membrane-spanning β-pore upon binding to claudin-4 (**Fig. 3B**). This model shows that COPs are sterically occluded from accessing their surface epitopes on cCpE due to CpE oligomeric assembly and N-terminal β-pore engagement (**Fig. 3C**). Based on this model we hypothesize that both COP-2 and COP-3 would be ineffective at preventing CpE-induced cytotoxicity. We test this hypothesis using a cell-based cytotoxicity assay and show that cell media supplemented with COPs do not prevent nor alter cytotoxicity induced by CpE (**Fig. 3C**). This finding verifies our hypothesis and provides evidence that our model of the CpE β-pore is potentially useful to predict the functional effects of structural changes to CpE upon binding claudin-4 and forming a cytotoxic β-pore.

Based on these data and results we propose a framework for using COP-like molecules to target and obstruct CpE cytotoxicity (**Fig. 4**). We envision three Strategies based on the proposed sequence of events that lead to CpE β-pore formation (17, 18). These include developing COPs that: 1) obstruct formation of claudin-4/CpE “small complexes”; 2) stabilize the native fold of CpE’s N-terminus to prevent its structural rearrangement that induces cytotoxicity via β-pore formation; and 3) obstruct CpE oligomerization prior to β-pore assembly. Using our models for the “small” and claudin-4-bound CpE β-pore complexes we identify five areas where targeted binding by COP-like sFabs could obstruct CpE binding, conformational changes, or oligomerization, potentially preventing CpE-based cytotoxicity. Area A is a surface pocket between β8 and β9 on the cCpE domain of CpE (**Fig. 4A**); and Area B comprises the two ECS of claudins and includes the NPLVA^153^ motif that imparts high-affinity CpE binding (**Fig. 4B**) (23). Development of COPs that bind Area A or B would inhibit formation of the “small complex” by obstructing areas known to facilitate claudin-4/cCpE interactions, thus preventing CpE pre-pore and β-pore complex formation (Strategy 1). Area C is a region of CpE’s N-terminus that we hypothesize structurally rearranges upon CpE assembly, which, like lysenin, then forms a long, antiparallel β-barrel to create the membrane-penetrating β-pore (**Figs. 3B** and **4C**). Discovery of COPs that bind Area C may stabilize CpE’s N-terminus to disable its β-pore-forming conformational transition (Strategy 2)—they may also obstruct CpE oligomerization (Strategy 3). Finally, Area D includes strand β1 and helix α1 of cCpE; while Area E comprises two loops that connect β2 to β3 and β8 to β9 (**Fig. 4C**). Based on our models, Areas D and E also represent parts of cCpE where COP binding would sterically obstruct CpE oligomerization. In summary, our models reveal that: 1) for COP binding to Areas A or B to be effective at obstructing cytotoxicity they will need to be present before or during CpE production to prevent “small complex” formation; and 2) COP binding to Areas C, D, or E would not prevent “small complex” formation and thus could be effective post-production of CpE but would need to be present prior to CpE’s oligomeric assembly of its β-pore. This proposed structure-based framework can inform therapeutic strategies that utilize unyet developed COPs to target CpE, claudin-4, or claudin-4/CpE complexes. COPs with traits that enable recognition of the abovementioned Areas could obstruct CpE β-pore formation and thus prevent CpE-linked cytotoxicity. Such molecules would be potent therapeutics, capable of treating gastrointestinal illnesses linked to CpE that annually afflict millions of humans and domesticated animals globally, resulting in large economic burdens and losses in quality of life.

**Figure 4.**
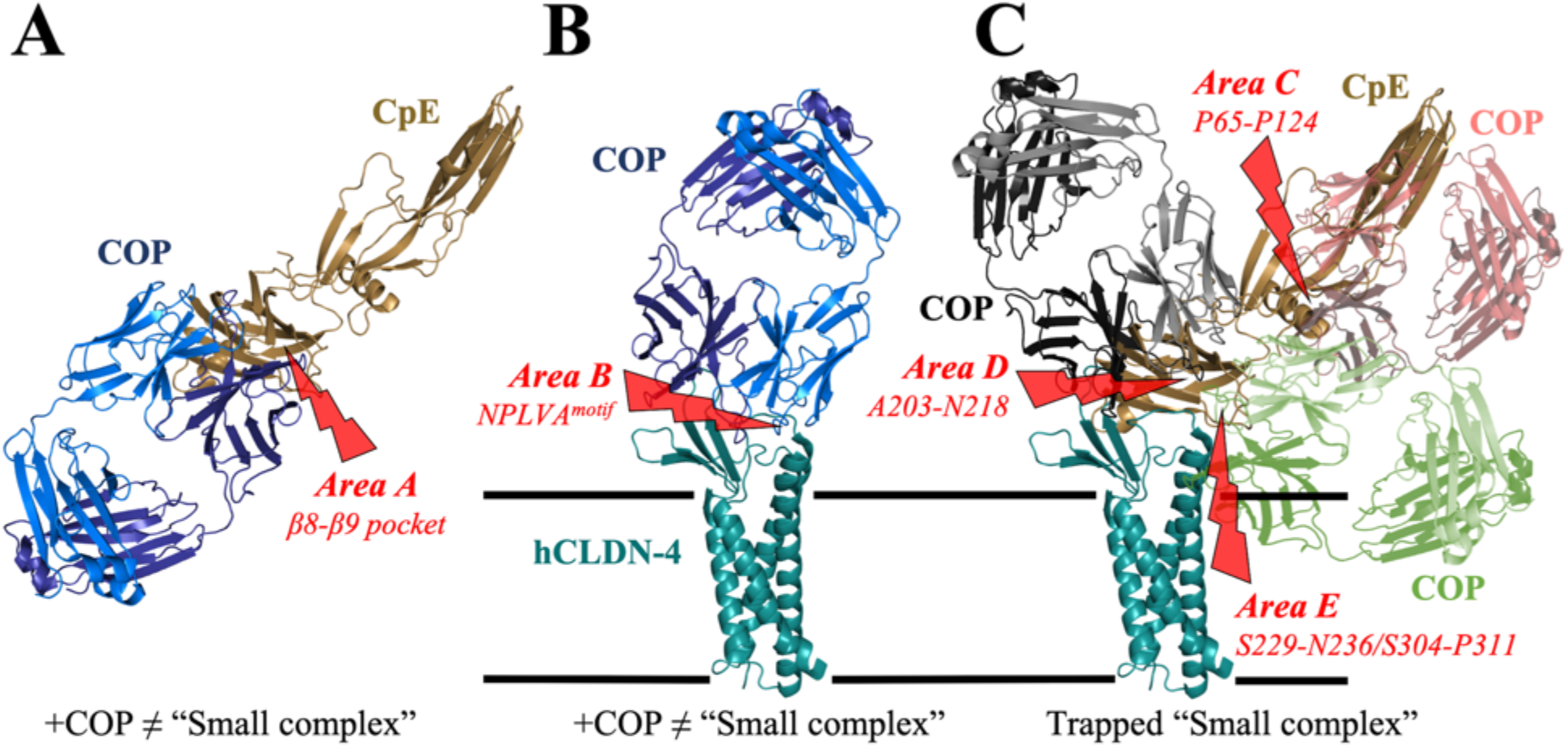
COP Targeting Approaches to Obstruct CpE Cytotoxicity. (**A**) COPs (blue) that target the cCpE domain of CpE (copper) by binding to *Area A*, a solvent accessible pocket between *β*8 and *β*9 of cCpE that is known to drive high-affinity binding to claudin-4’s NPLVA^153^ motif, which is critical for “small complex” formation. (**B**) COPs (blue) that target human claudin-4 (hCLDN-4, teal) by binding to *Area B*, the two ECS of claudins, which both interact with cCpE to initiate “small complex” formation. (**C**) COPs (red, black, green) that target CpE (copper) by binding to *Areas C*, *D*, and *E*. *Area C* comprises residues Pro65-Pro124 of CpE’s N-terminus, which contains a purported Thr92-Gly105 α-helix used for β-pore formation. *Area D* comprises residues Ala203-Asn218 of cCpE, which contains its *β*1 strand and α1 helix. *Area E* comprises residues Ser229-Asn236 and Ser304-Pro311 of cCpE, which contain two loops that connect strands *β*2 to *β*3 and *β*8 to *β*9.

In this study we progress a novel workflow for rapid determination of claudin/enterotoxin structures using practical 200 keV cryoEM instrumentation that yields 4-7 Å resolutions. The approach appears amenable to other claudins in a variety of membrane mimetics, and is enabled by 50 kDa COPs that add mass, rigidity, and act as fiducial marks for cryoEM (***SI Appendix*, Fig. S10**). The COPs are versatile, capable of being used for cryoEM and/or as crystallization chaperones for X-ray crystallography. Although the cryoEM workflow described here produces modest resolution structures, it must be considered that these complexes are small by cryoEM standards at 35 kDa. We estimate that their already structurally detailed maps could be improved to 3-4 Å using 300 keV microscopes (***SI Appendix*, Fig. S5**). Achieving these resolutions would equal current crystallography results for claudin/cCpE complexes, and have further advantages like faster structural throughput timeframes and bypassing crystallization and diffraction bottlenecks. Thus, COPs could be employed as tools to rapidly expand structural knowledge of other claudin orthologs or paralogs that bind enterotoxins. In this and their potential obstructing capacities, COPs and COP-like sFabs may provide new insights useful for developing treatments for CpE-based diseases or to aid design of novel cCpE- and CpE-based therapeutics that modulate TJ barriers.

## METHODS

### Claudin-4 and Enterotoxin Expression and Purification

Methods followed those described previously (22, 23). Briefly, claudin-4, cCpE, and CpE with C-terminal decahistidine tags preceded by thrombin cleavage sites (claudin-4-His_10_, cCpE-His_10_, and CpE-His_10_) were cloned into pFastBac1 (ThermoFisher) and expressed in Tn5 (*Trichoplusia ni*, High Five, Expression Systems, LLC). Cell pellets resuspended in Lysis buffer (50 mM Tris pH 8.0, 150 mM NaCl, 1 mM PMSF, and EDTA-free SigmaFast protease tablets (Sigma)) were sonicated, supplemented with 1 M NaCl, then ultracentrifugation at 100,000 xg for 1 hour. For enterotoxins, the supernatant was saved, 15 mM imidazole was added along with NiNTA resin, and the solution was incubated for 12 hours at 4°C. For claudin-4, the supernatant was removed, and the membrane pellet was resuspended in Lysis buffer and the protein was solubilized with 1% (w/v) n-dodecyl-β-D-maltopyranoside (DDM, Anatrace) and 0.04% cholesteryl hemisuccinate (CHS, Anatrace) overnight at 4°C. Insoluble protein was removed by ultracentrifugation at 100,000 xg for 30 minutes, and the supernatant was treated with 15 mM imidazole along with NiNTA resin, and the solution was incubated for 12 hours at 4°C. For claudin-4, the bound protein was captured and washed with 5 column volumes of Buffer A (50 mM Tris pH 7.4, 500 mM NaCl, 25 mM imidazole, and 0.087% DDM) and B (Buffer A containing 300 mM NaCl and 40 mM imidazole). Buffer T (50 mM Tris pH 8.0, 150 mM NaCl and 0.04% DDM) was used to release and capture proteins from the resin after treatment with thrombin. For enterotoxins, purification was similar to claudin-4 except DDM was not added to buffers. These proteins were then used for sFab panning or biochemical, biophysical, and structural analyses. For enterotoxins used for binding studies, the proteins were eluted off of NiNTA using Elution buffer (Buffer T containing 400 mM imidazole) to keep the His_10_ tag. Eluted enterotoxin-His_10_ was dialyzed in size-exclusion chromatography (SEC) buffer (10 mM Hepes pH 7.4, 100 mM NaCl, and 4% glycerol), concentrated to 1 mg/mL, then flash-frozen in liquid nitrogen and stored at −80°C until use.

### Generation and Validation of COPs Using Phage Display

DDM-solubilized claudin-4 was biotinylated using N-hydroxysuccinimide polyethylene glycol biotin (NHS-PEG4-biotin, ThermoFisher) by mixing 5.6 μM claudin-4 with 16.8 μM NHS-PEG4-biotin, followed by incubation on ice for 2 hours. Excess cCpE with the His_10_ removed was added at a ratio of 1:1.5 (moles:moles), then free biotin and unbound cCpE was removed by loading the sample onto a Superdex 200 Increase 10/300 GL (Cytiva) equilibrated in SEC buffer containing 0.04% DDM. The sFab panning was performed using the sFab library E (33, 34). Binding was assayed in Selection buffer (25 mM Hepes pH 7.4, 150 mM NaCl, and 1% bovine serum albumin). A first round of panning was performed manually using 200 nM of biotinylated claudin-4/cCpE in DDM immobilized onto magnetic beads, and following three washes with Selection buffer, the beads enriched for phage expressing claudin-4/cCpE-specific sFabs were used to infect log-phase *E. coli* XL1-Blue cells. Phages were amplified overnight in 2xYT media supplemented with 100 μg/ml ampicillin and M13-KO7 helper phage (10^9^ pfu/ml). Selection stringency was then increased by four additional rounds of panning using decreasing claudin-4/cCpE concentrations down to ~20 nM. For each round, the amplified phage pool from each preceding round was used as the input. For rounds two to five, panning was performed semi-automatically using a Kingfisher magnetic beads handler (ThermoFisher). Non-specific binding of sFabs to detergent was reduced by using >0.87% DDM in later rounds. Bound phage particles were removed by elution from beads using 1% Fos-choline-12.

The initial validation was performed by single-point phage ELISA using individual clones from later selection rounds. ELISAs were performed in 96-well plates (Nunc) coated with 2 μg/ml neutravidin and blocked with Selection buffer. *E. coli* XL1-Blue colonies containing phagemids were used to inoculate 400 μl 2xYT media containing 100 μg/ml ampicillin and 10^9^ pfu/ml M13-KO7 helper phage. Phages were amplified overnight in 96-well deep-well blocks at 37°C with shaking at 280 rpm. Phages were then diluted 1:10 into Selection buffer and assayed against claudin-4/cCpE in DDM or buffer containing only DDM micelles. Biotinylated claudin-4/cCpE was immobilized at room temperature for 30 min then incubated with phage dilutions. Bound phage were detected with TMB substrate (ThermoFisher) following a 30 minutes incubation with horseradish peroxidase-conjugated anti-M13 monoclonal antibody (GE Healthcare). Absorbance was measured at 450 nm after quenching the reaction with 1.0 M HCl. Wells containing 1% DDM were used to detect non-specific binding.

### COP Expression and Purification

Two sFabs termed <u>C</u>pE <u>O</u>bstructing <u>P</u>rotein-2 and −3 (COPs), from phage ELISA were selected and sequenced at the University of Chicago Comprehensive Cancer Center DNA Sequencing facility. Unique clones for each COP were sub-cloned in pRH2.2 using the In-Fusion Cloning kit (Takara Bio). Sequence-verified COPs were transformed into *E. coli* BL21-Gold cells (Agilent) and then used to inoculate overnight cultures. The inoculates were then used to seed 1 L of 2xYT media containing 100 μg/ml ampicillin. Cultures were grown to an OD_600_ of 0.8, induced for 4 hours at 37°C, then cells were harvested using centrifugation. Cell pellets were resuspended in COP Lysis buffer (20 mM Hepes pH 7.4, 150 mM NaCl, 0.5 mM MgCl_2_, 1 mM PMSF, and 1 μg/ml DNase I). Cells were sonicated and lysates were incubated at 60°C for 30 minutes to remove proteolyzed fragments. Samples were cooled rapidly on ice then cleared by centrifugation. Supernatants were filtered by 0.45 μm and loaded onto a 5 mL HiTrap MabSelect SuRe column (GE Healthcare) equilibrated with COP Wash buffer (20 mM Hepes pH 7.4 and 500 mM NaCl). The column was washed with 10 column volumes of COP Wash buffer and COPs were eluted with 0.1 M acetic acid. Eluted COPs were loaded onto a 1 mL Resource S column (GE Healthcare) equilibrated with COP Buffer A (50 mM sodium acetate pH 5.0) and washed with 10 column volumes of this buffer. The COPs were eluted by linear 0–50% gradient with COP Buffer B (COP Buffer A containing 2 M NaCl). COP-containing fractions were pooled and dialyzed overnight at 4°C in SEC buffer.

### Biochemical Characterization of COP Binding

We used post-NiNTA purified un-tagged claudin-4 and enterotoxins, and post-affinity purified COPs for these analyses. For claudin-4/COP and enterotoxin/COP studies, 50 μg enterotoxins was used and excess COPs were added at a 1:1.2 molar ratio, incubated at room temperature for 1 hour, concentrated, 0.2 μm filtered, then loaded onto a Superdex 200 column equilibrated in SEC buffer containing 0.04% DDM. For claudin-4/enterotoxin/COP studies, 50 μg claudin-4 was used and excess enterotoxins were added at a 1:1.2 molar ratio, then COPs were added at a 1:1 molar ratio to enterotoxins. Complexes were incubated, concentrated and filtered, and loaded onto a Superdex200 column equilibrated in SEC buffer and 0.04% DDM. Complex formation was assessed by observed decreases in the elution times of the uncomplexed peak fractions. Peak fractions containing complexes were pooled, unboiled, and evaluated for the presence of each protein by SDS-PAGE using 4-20% agarose gradient gels.

### Biophysical Characterization of COP Binding Using Bio-layer Interferometry (BLI)

BLI analyses were performed at 25°C at an acquisition rate of 5 Hz averaged by 20 using an Octet© BLItz System (FortéBio/Sartorius) with assays were designed and setup using Blitz Pro 1.3 Software. A typical experiment consisted of the following steps: sensor equilibration (30 seconds), protein loading (200 seconds), baseline (60 seconds), and association and dissociation (300 seconds each). For the loading step, 4 μl of proteins were loaded in the drop holder, while all other steps were performed in transparent 600 μl microtubes using 250 μl sample volumes. All measurements were performed in SEC buffer containing 0.04% DDM. For enterotoxin/COP studies, 5 μM (70 μg/ml) of wild type or mutant cCpE-His_10_ or CpE-His_10_ were loaded on NiNTA (Dip and Read^™^) sensors then dipped into a 0-1 μM range of four concentrations of COPs for the association steps. For claudin-4/enterotoxin/COP studies, 5 μM DDM-solubilized claudin-4 biotinylated with NHS-PEG_4_-biotin as before was pre-complexed with excess wild type or mutant enterotoxins without a His_10_ at a ratio of 1:2 (moles:moles) for 1 hour at room temperature. Claudin-4/enterotoxin complexes were loaded on Streptavidin-SA (Dip and Read^™^) sensors and the measurements were performed as above using 0-10 μM range of concentrations depending on the analyte. For all studies, measurements were repeated in at least duplicate, and the time courses for association and dissociation were fit to one-site binding model using the BLItz Pro 1.3 Software. No significant nonspecific binding of COPs to unloaded NiNTA or streptavidin sensors were detected.

### CryoEM Sample Preparation

DDM-solubilized claudin-4 was exchanged into 2,2-didecylpropane-1,3-bis-β-D-maltopyranoside (LMNG, Anatrace) or amphipol A8-35 (Anatrace). For exchange into LMNG the protein bound to NiNTA resin was sequentially washed with Buffer A, B, and T containing 0.087, 0.087, and 0.04% LMNG, respectively, before releasing the bound protein from resin via thrombin cleavage. To prepare sample in amphipol A8-35, 1 mg of post-thrombin digested claudin-4 in DDM was treated with 4 mg of amphipol (1:4 w/w) for 2 hours before removing detergent via addition of 400 mg SM-2 biobeads (Bio-Rad). Excess cCpE was added to each claudin-4 sample at a molar ratio of 1:1.5, incubated at room temperature for 1 hour, then excess COPs were added at a 1.5:1 ratio to cCpE. After 1 hour at room temperature each sample was concentrated, 0.2 μm filtered, then loaded onto a Superdex 200 Increase 10/300 GL equilibrated in SEC buffer without glycerol but with 0.003% LMNG or no detergent for amphipol samples. Peak fractions from SEC containing claudin-4/cCpE/COP complexes were collected and concentrated to 6-8 mg/mL for use in cryoEM analyses.

Grids for cryo-EM analyses were prepared using a Vitrobot Mark IV (ThermoFisher) plunge freezing apparatus. Aliquots (3 μL) of claudin-4/cCpE/COP complexes in LMNG or amphipol were applied to Quantifoil R1.2/1.3 200 mesh grids that were glow-discharged for 45 seconds at 15 mA in a Pelco easiGlow (Ted Pella Inc) instrument. Protein solutions were applied to grids at 4°C and 100% relative humidity and allowed to adsorb on the grid for ~30 seconds before blotting. Grids were blotted for 5-8 seconds with a blot force of 1, and plunge frozen into liquid ethane cooled by liquid nitrogen. Grids were stored in liquid nitrogen prior to imaging.

### CryoEM Data Collection and Processing

CryoEM data collection was performed on a Talos Arctica (ThermoFisher) equipped with a Falcon III (ThermoFisher) direct electron detector at Michigan State University. Grids were screened for thin ice and good particle distribution, and data collection was performed using EPU software (ThermoFisher). Movies of claudin-4/cCpE/COP-2 in LMNG were collected on the Falcon III detector in counting mode at 92,000x magnification with a pixel size of 1.12 Å, a defocus range of 0.8 to 2.6 μm, and a total dose of ~32 electron/Å^2^ fractionated over 51 total frames. Movies of claudin-4/cCpE/COP-3 in amphipol were collected at 120,000x magnification with a pixel size of 0.87 Å, a defocus range of −1 to −2.2 μm, and a total dose of ~40 electron/Å^2^ fractionated over 42 total frames.

All micrograph and particle processing was performed in CryoSPARC (40). Patch-motion correction and patch-CTF correction were used to correct for beam-induced motion and calculate CTF parameters from the motion-corrected micrographs. Blob-based template picking followed by 2D classification was used to generate templates that were subsequently used for template-based particle picking. Particles identified from this template-based picking procedure were subjected to several rounds of 2D classification, followed by *ab initio* 3D reconstruction, heterogeneous refinement, and non-uniform refinement. In the case of the COP-3 complex, local refinement using a soft mask around cCpE and COP-3 was also used to improve resolution at the complex’s interface.

### CryoEM Model Building, Refinement, and Structure Determination

Using post-processing maps we first built the structure of claudin-4/cCpE/COP-2 using the 6.9 Å map and the crystal structure of claudin-4 in complex with cCpE (PDB ID 7KP4) (23). 7PK4 was manually docked by placing the four TMs of claudin-4 into density present in the LMNG micelle using Coot (41). TM placement was validated using density corresponding to bulky side chains, like Trp18 in TM1 of claudin-4. Initial placement of the TM region showed that only minor alterations to claudin-4’s ECS and cCpE would be required to fit well within the cryoEM map. For this part of the structure, the map volume allowed placement of most claudin-4 and cCpE residues, with the exception of claudin-4’s C-terminus and cCpE’s N-terminus. We next built COP2 by first using a high-resolution crystal structure of a sFab (PDB ID 6CBV). COP-2 was manually docked into the additional cryoEM density using Coot, and then divergent residues of COP-2 were manually mutating from the 6CBV template using sequence alignments as a guide (42). Once the three proteins were fit manually, each of the four protein chains were rigid body refined in Coot to place the structure within the cryoEM map volume. Model building was done using Coot and once complete, the structure was further refined using a combination of molecular dynamics flexible fitting simulations using Namdinator followed by real-space refinement of the model into the cryoEM density map using Phenix phenix.real_space_refine (43, 44). The final structural model required secondary structure and Ramachandran restraints to optimize the model-to-map fit and overall geometry. ***SI Appendix*, Table S1** shows data collection, refinement, and validation statistics for the claudin-4/cCpE/COP-2 structure.

The structure of claudin-4/cCpE/COP-3 was determined using both a masked and unmasked strategy, which produced *focused* and *whole* maps that resolved to 3.8 and 5.0 Å, respectively. A mask was applied to COP3 in the complex in the former case while all three proteins were included in the latter. The final model of the claudin-4/cCpE/COP-2 complex was manually docked then COP-3 was made from COP-2 by mutating divergent residues using sequence alignments. Each protein chain in the complex was then individually refined as rigid bodies into the cryoEM maps using Coot. While the *focused* map had added features for COP-3 it lacked features for claudin-4 and cCpE such that refinements resulted in poor model-to-map fits. Thus, all three proteins were built and fit manually within the *whole* map volume using Coot and the final model was refined using Namdinator and Phenix phenix.real_space_refine using secondary structure and Ramachandran restraints (43, 44). ***SI Appendix*, Table S1** shows data collection, refinement, and validation statistics for the claudin-4/cCpE/COP-3 structure.

The programs used to visualize and build the structures included Coot, PyMOL, and Chimera, refined using Phenix, and figures were made using PyMOL—using the SBGrid Consortium Software Suite (41, 45–48).

### Mutagenesis of cCpE

The pFastBac1 cloned cCpE-His_10_ was altered using site-directed mutagenesis. Mutants were generated with the following forward primers and their equivalent reverse compliments:

cCpE^1^ 5’ -gcgcggatccgccaccgcatcaacggacattatgaaagaaatcctcgac-3’;
cCpE^2^ 5’ -ccatgtcaacggacattgaaaaagaagccgccgccttagctgctgcaacagaacgc-3’;
cCpE^2L^ 5’ -ccatgtcaacggacattgaa<u>aaa</u>gaagccgccgccgccgctgctgcaacagaacgc-3’;
cCpE^3^ 5’ -gttgactttaacatttactccaacgccgccgctgcccttgtcaaactcgaacaatcgctc-3’;
cCpE^4^ 5’-catttactccaacaacttcaataacgctgccaaagccgcacaatcgctcggagatggtg-3’. Protocols for expression and purification of all cCpE-His_10_ mutants were identical to those of wild type cCpE-His_10_.

### Structural Modeling of Claudin-bound CpE Pore

A model of claudin-4/CpE was made by superposing CpE (PDB ID 3AM2) onto cCpE from our cryoEM structure of the COP-2 complex, making a claudin-4/CpE/COP model. To approximate the CpE β-pore, we modeled it after lysenin, a β-pore toxin with homology to CpE, using an available cryoEM structure (PDB ID 5GAQ) (37, 49). Using a crystal structure of the lysenin monomer (PDB ID 3ZXD), we superposed 3ZXD onto one monomer of the 5GAQ nonamer, then superposed the CpE portion of our claudin-4/CpE/COP model onto 3ZXD (50). We next removed 3ZXD. This process integrated claudin-4/CpE/COP into lysenin, making the model for the claudin-4-bound CpE β-pore. These models were used to predict COP influence on CpE-induced cytotoxicity.

### Cell-Based Cytotoxicity Assay

Recombinant baculoviruses containing claudin-4-His_10_ were produced using established methods and the cytotoxicity assay was performed using methods previously described (23). To 18 wells of a 24-well cell culture plate, 0.5 mL of 1.0×10^6^ adherent Sf9 (*Spodoptera frugiperda*, Expression Systems) cells were added. After 1 hour at 27°C, virus containing claudin-4-His_10_ was added at a MOI of 1.0. Cultures were rocked gently then placed at 27°C for 48 hours. The 18 wells were divided into three, 6-well groups. To duplicate wells of group one and three, 22 μg (0.9 μM) of COP-2 was added; to a second pair of wells, 22 μg (0.9 μM) of COP-3 was added; and to the third pair of wells no COP was added. To duplicate wells of group two, 100 μg (4 μM) of COP-2 was added; to a second pair of wells, 100 μg (4 μM) of COP-3 was added; and to the third pair of wells no COP was added. For group one, nothing more was added; this group contained only cells expressing claudin-4-His_10_. For group two, 25 μg (3.3 μM) purified cCpE in SEC buffer was added to the culture medium. For group three, 12.5 μg (0.7 μM) CpE in SEC buffer was added to the culture medium. The COP amounts added correspond to 1.2-fold molar excess of COP to enterotoxin. After addition of COPs and/or enterotoxins, Sf9 cells were placed at 27°C for 12 hours, then CpE cytotoxicity was quantified using a cell viability analysis. This was done by gently removing 250 μL of Sf9 cells from each well, centrifuging at 200 xg for 2 minutes, removing 200 μL of media, then adding 50 μL of 0.04% trypan blue. After 5 minutes, 10 μL of stained cells were transferred to a Countess cell counting chamber slides and counted automatically using a Countess 3 automated cell counter (ThermoFisher). Each well was counted two times independently (representing four readings per sample, 36 for nine samples). Viability counts consisted of dividing the total number of live cells (unstained) by the total number of cells. Average cell viability for enterotoxin treated cells were compared to Sf9 cells expressing claudin-4 that were not treated with either enterotoxins or COPs. Data was plotted as a mean with standard deviation in GraphPad Prism 9 for macOS.

## ACKNOWLEDGMENTS

Research reported in this publication was supported by the National Institute of General Medical Sciences of the National Institutes of Health under Award Number R35GM138368 (to A.J.V.) and R01GM117372 (to A.A.K). The content is solely the responsibility of the authors and does not necessarily represent the official views of the National Institutes of Health. Also, we are grateful to the RTSF Cryo-Electron Microscopy Core at Michigan State University and Dr. Sundarraman Subramanian for providing time and support for cryoEM data collection.

## AUTHOR CONTRIBUTIONS

A.J.V. designed the research; performed claudin and enterotoxin expression, purification and sample preparation for structure/function studies, biochemical characterization, cytotoxicity assays; determined cryoEM structures; analyzed data; made figures; and wrote the paper; B.J.O. performed the cryoEM, determined cryoEM structures, and edited the paper; P.K.D. performed sFab phage display screening and validation; P.K.D. and S.K.E. performed COP expression and purification; S.R. performed binding measurements; S.R. and C.O. performed claudin and enterotoxin expression and purification for binding studies; A.A.K. developed the sFab phage display library, designed the sFab-related research, and edited the paper.

## COMPETING INTERESTS

The authors declare no competing interests.

## DATA AVAILABILITY

The cryoEM structure of claudin-4/cCpE/COP-2 has accession code 7DTM in the Protein Data Bank (PDB) and cryoEM maps of this complex have been deposited to the Electron Microscopy Data Bank under accession code EMD-25834. The cryoEM structure of claudin-4/cCpE/COP-3 has PDB accession code 7DTM and cryoEM maps of this complex have been deposited to the Electron Microscopy Data Bank under accession code EMD-25835 (*whole*) and EMD-25836 (*focused*).

## Supplementary Information (SI) Appendix

**SI Figure S1.**
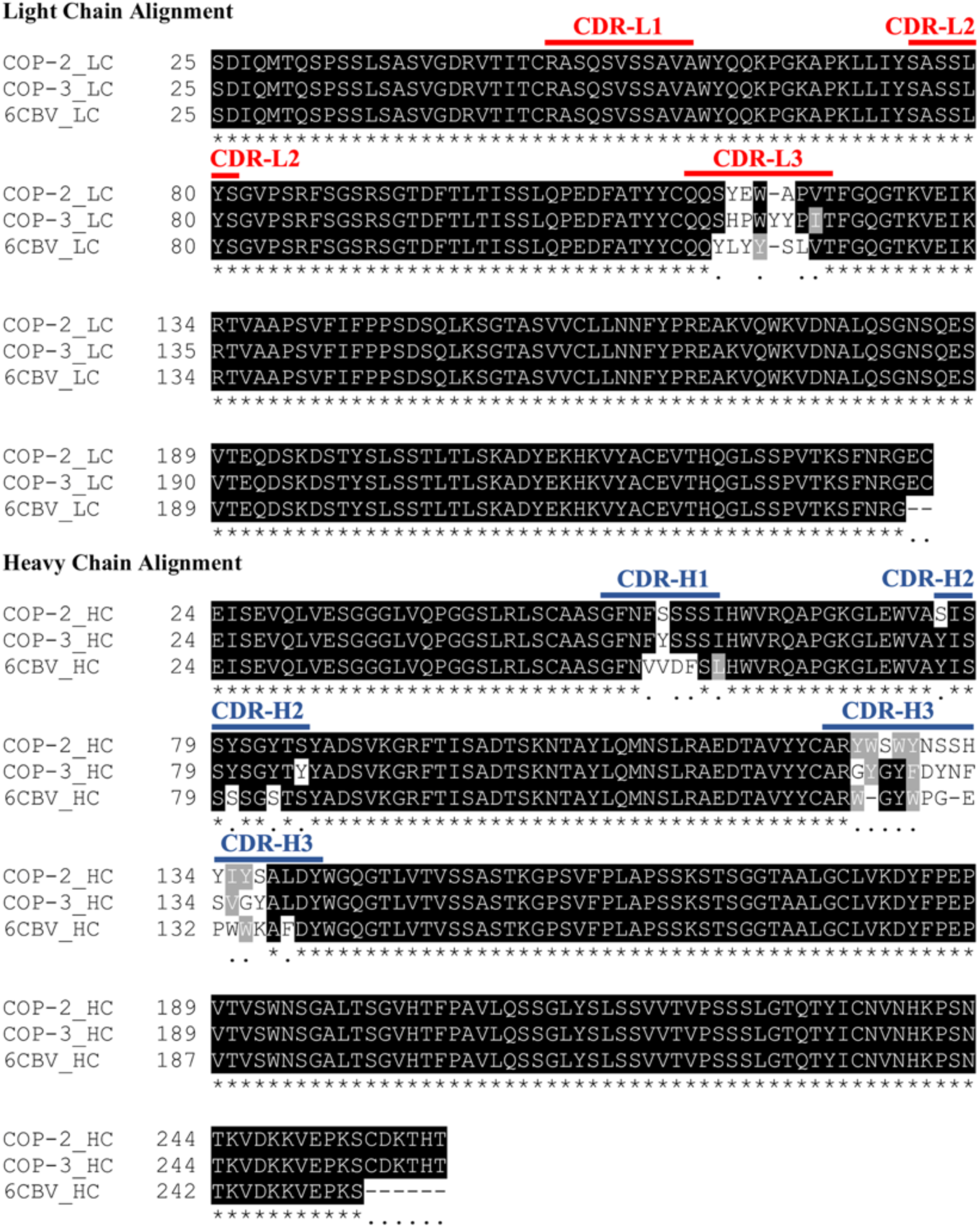
Sequence Alignment of COPs. The L and H chains of COP-2 and COP-3 were sequenced and aligned against the sequence of a generic sFab, PDB ID 6CBV, using T-Coffee (51). Highlighted in the sequences are the the three CDRs from the L chain (red) and three CDRs from the H chain (blue).

**SI Figure S2.**
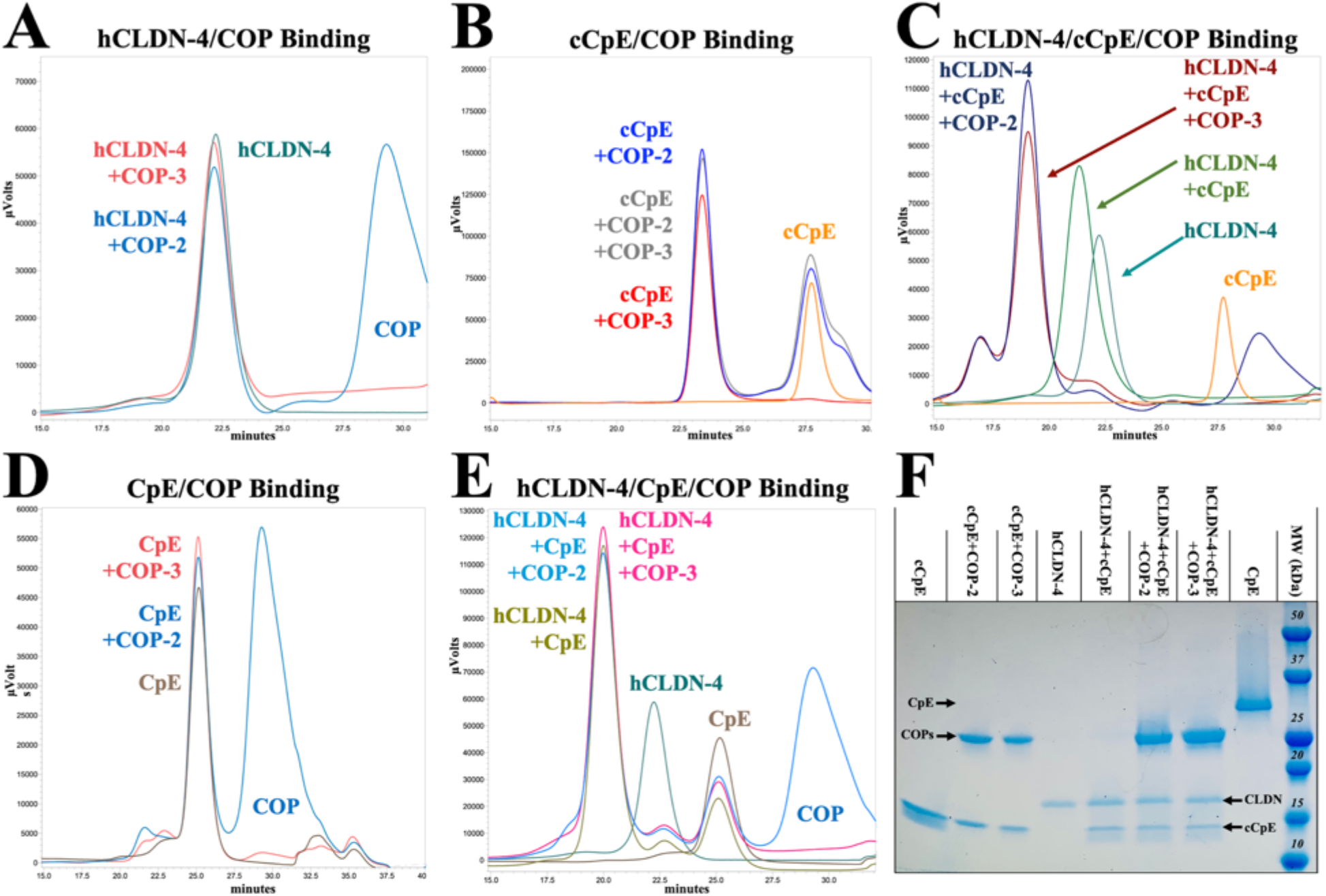
Biochemical Characterization of COP Molecular Recognition. Human claudin-4 (hCLDN-4), cCpE, CpE, and COP-2 or COP-3 were incubated together and then injected onto a SEC column equilibrated in DDM and monitoring using 280 nm absorbance. The SEC traces depict: COP-2 (blue) and/or COP-3 (red) binding to (**A**) claudin-4 alone (green); (**B**) cCpE alone (orange); (**C**) claudin-4/cCpE complexes (green); (**D**) CpE alone (brown); and (**E**) claudin-4/CpE complexes (brown green). (**F**) SEC peak fraction from **A-E** were pooled and subjected to SDS-PAGE.

**SI Figure S3.**
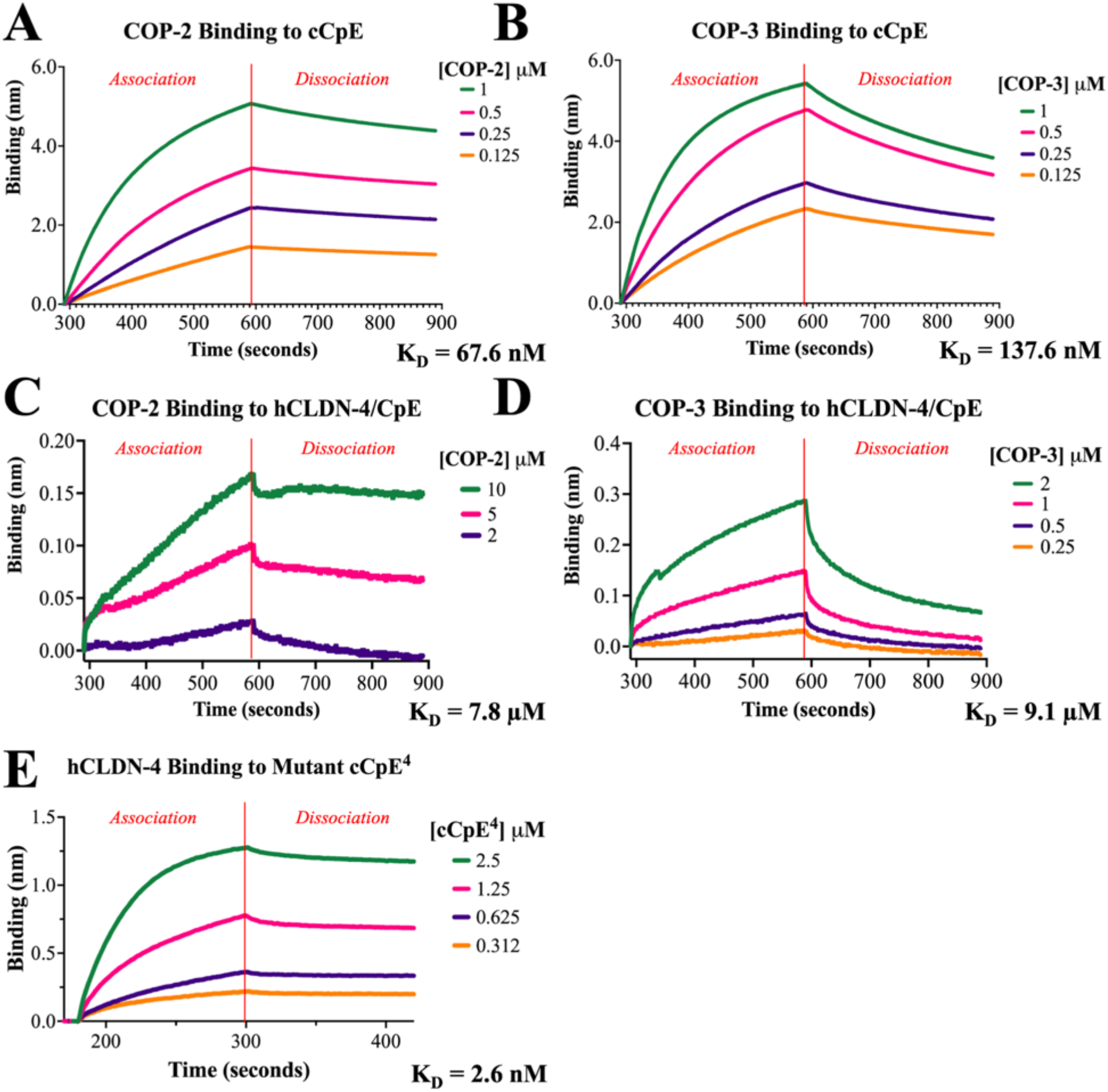
Biophysical Binding Measurements Using BLI. (**A**) COP-2 and (**B**) COP-3 binding to cCpE-His_10_ immobilized on NiNTA biosensors. (**C**) COP-2 and (**D**) COP-3 binding to biotinylated human claudin-4 (hCLDN-4) in complex with cCpE immobilized on streptavidin (SA) biosensors. (**E**) Mutant cCpE^4^ binding to biotinylated hCLDN-4 immobilized on SA biosensors. Analyte concentrations are colored from high to low (green, magenta, purple, orange) and a red solid line depicts the transition from association to dissociation phase. BLI sensorgrams collected using BLItz Pro 1.3 Software were re-plotted in Prism 9. Sensorgrams represent a single experiment set from duplicate measurements.

**SI Figure S4.**
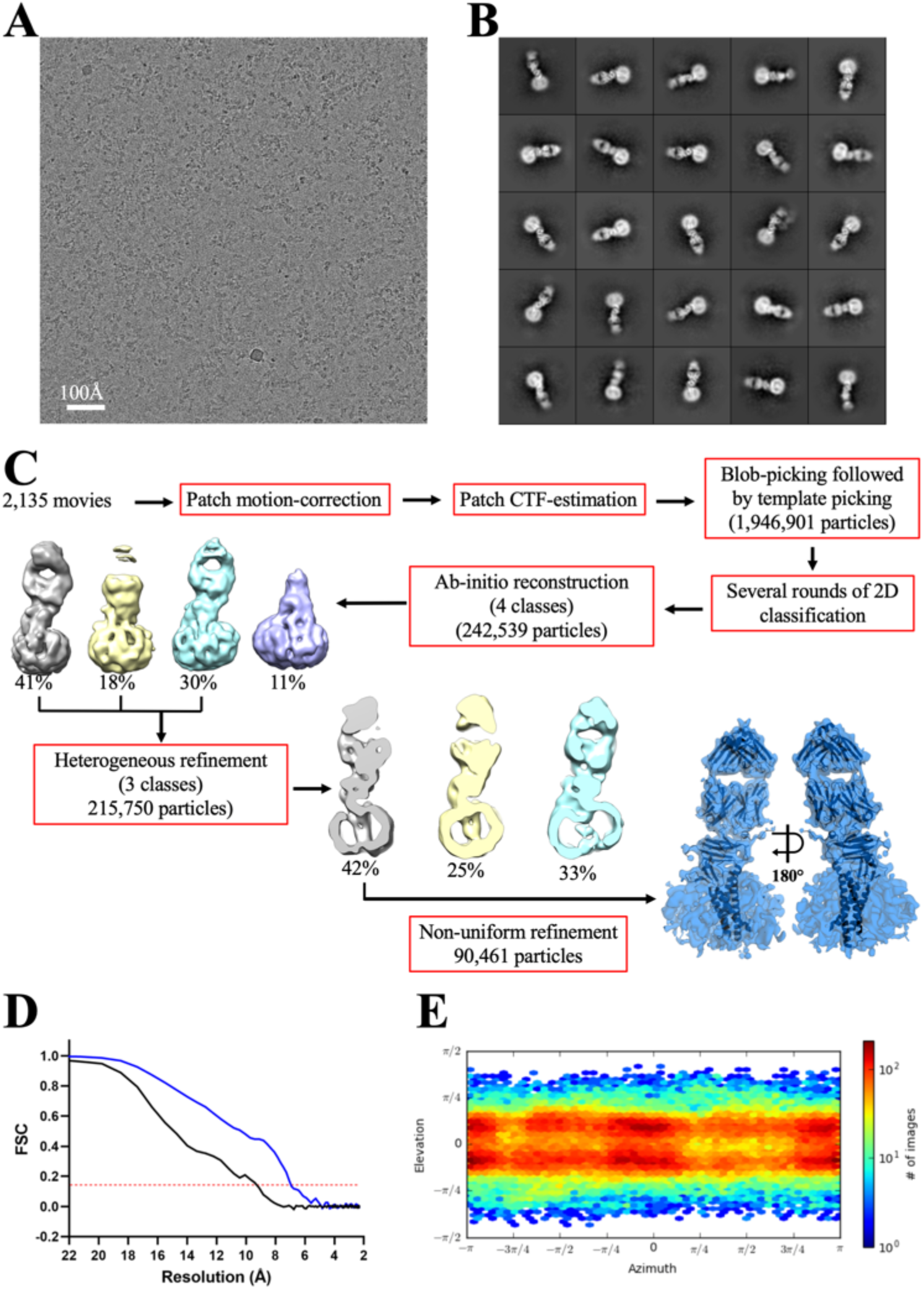
CryoEM Data Processing for Claudin-4/cCpE/COP-2 Complexes. (**A**) Electron micrograph showing the distribution of claudin-4/cCpE/COP-2 complexes in ice. (**B**) Representative 2D class averages of claudin-4/cCpE/COP-2 complexes showing the clear binding of the COP-2 sFab, and signal for TMs within the LMNG detergent micelle. (**C**) CryoEM data processing workflow. (**D**) Fourier Shell Correlation (FSC) curves from gold-standard refinement. The 0.143 FSC cutoff is indicated by a dashed red line. The black line indicates FSC without a mask applied, and the blue line indicates FSC with a tight mask applied around the protein and detergent micelle. (**E**) Angular distribution of particles in the final refinement, demonstrating an absence of pure top views.

**SI Table S1.** CryoEM Data Collection, Refinement and Validation Statistics.

|  | Claudin-4/cCpE/<br>COP-2<br>EMDB-25834<br>PDB ID 7DTM | Claudin-4/cCpE/<br>COP-3<br>EMDB-25835 (*whole*)<br>PDB ID 7DTN | Claudin-4/cCpE/<br>COP-3<br>EMDB-25836 (*focused*) |
| --- | --- | --- | --- |
| **Data Collection and Processing** |  |  |  |
| Magnification | 92,000 | 120,000 | 120,000 |
| Voltage (keV) | 200 | 200 | 200 |
| Electron exposure (e-/Å<sup>2</sup>) | 32.0 | 39.6 | 39.6 |
| Defocus range (µm) | 0.8 - 2.6 | 1.0 - 2.2 | 1.0 - 2.2 |
| Pixel size (Å) | 1.12 | 0.871 | 0.871 |
| Symmetry imposed | C1 | C1 | C1 |
| Number of micrographs | 2,135 | 5,132 | 5,132 |
| Initial particle images (no.) | 1,946,901 | 3,758,631 | 3,758,631 |
| Final particle images (no.) | 90,461 | 305,927 | 305,927 |
| Map resolution (Å) | 6.9 | 5.0 | 3.8 |
| FSC threshold | 0.143 | 0.143 | 0.143 |
| **Refinement** |  |  |  |
| Initial model used (PDB ID) | 7KP4 | 7DTM | *N/A* |
| Model resolution (Å) | 6.1 | 4.5 | *N/A* |
| FSC threshold | 0.143 | 0.143 | *N/A* |
| Map sharpening *B* factor (Å<sup>2</sup>) | -500 | -150 | *N/A* |
| Model composition |  |  |  |
| Non-hydrogen atoms | 5695 | 5671 | *N/A* |
| Protein residues | 750 | 746 | *N/A* |
| *B* factors (Å<sup>2</sup>) |  |  |  |
| Protein | 164.03 | 413.26 | *N/A* |
| R.M.S. deviations |  |  |  |
| Bond lengths (Å) | 0.003 | 0.002 | *N/A* |
| Bond angles (°) | 0.726 | 0.681 | *N/A* |
| Validation |  |  |  |
| MolProbity score | 2.23 | 2.27 | *N/A* |
| Clashscore | 18.29 | 18.83 | *N/A* |
| Poor rotamers (%) | 0.00 | 0.48 | *N/A* |
| Ramachandran plot |  |  |  |
| Favored (%) | 92.58 | 91.73 | *N/A* |
| Allowed (%) | 7.42 | 8.27 | *N/A* |
| Disallowed (%) | 0.00 | 0.00 | *N/A* |

*N/A* = Not Applicable

**SI Figure S5.**
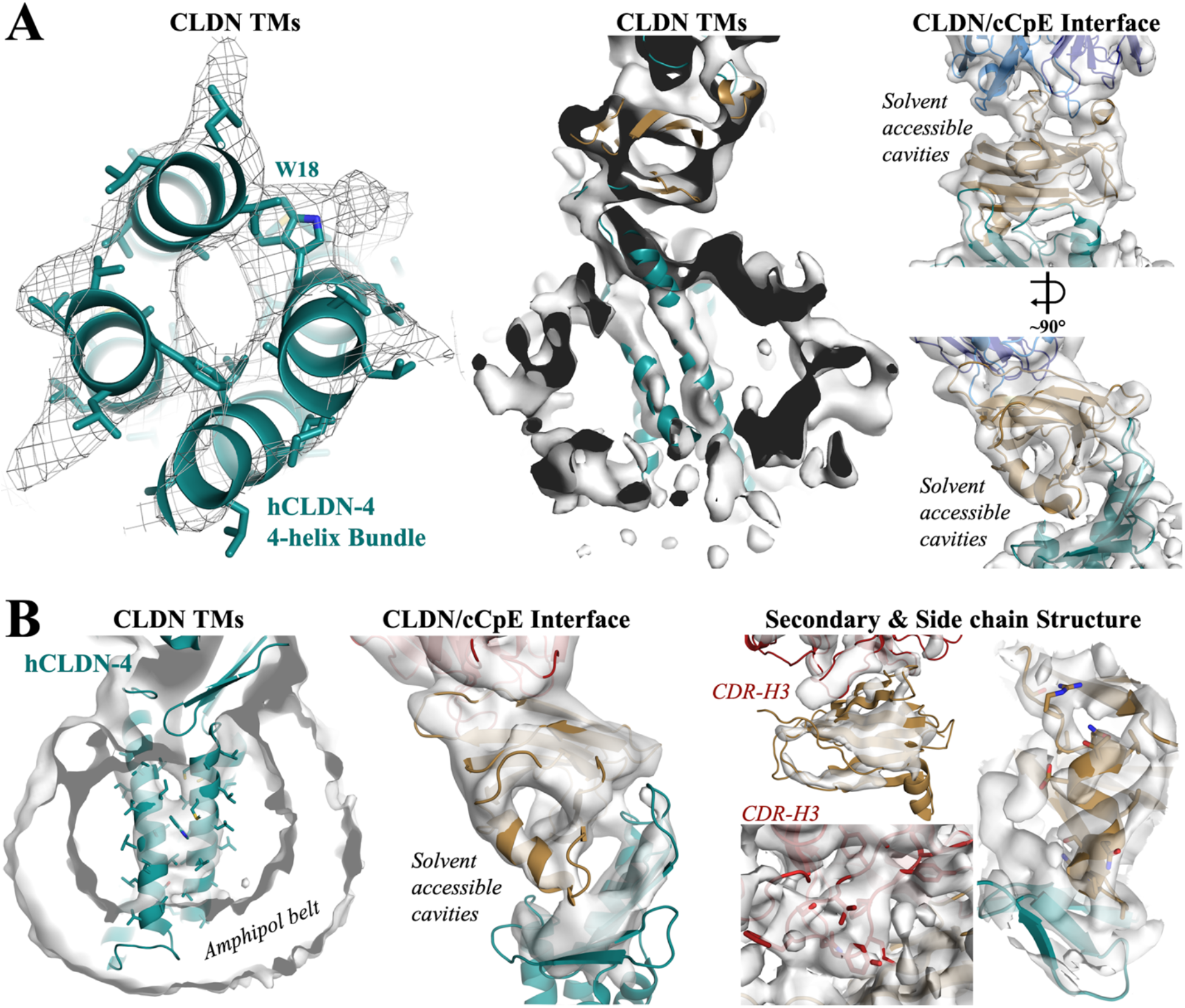
CryoEM Map and Structural Features. (**A**) Structure and corresponding 6.5 Å map from the human claudin-4 (hCLDN-4)/cCpE/COP-2 complex. Images depict: looking up through the TM bundle to the extracellular space with strong side chain density for Trp18 (left); a side view parallel to the membrane plane of individual TMs within the LMNG micelle (middle); and the claudin-4/cCpE and cCpE/COP-2 interfaces with various solvent accessible cavities resolved (right). (**B**) Structure and corresponding 4-4.5 Å map from the claudin-4/cCpE/COP-3 complex. Images depict: a side view parallel to the membrane plane of the TMs within the amphipol belt (left); claudin-4/cCpE interfaces with various solvent accessible cavities resolved (middle); and strand, helix, loop, and side chain density for aromatic amino acids at claudin-4/cCpE and cCpE/COP-3 interfaces (right). Proteins are shown as cartoons and colored as in **Fig. 1**.

**SI Figure S6.**
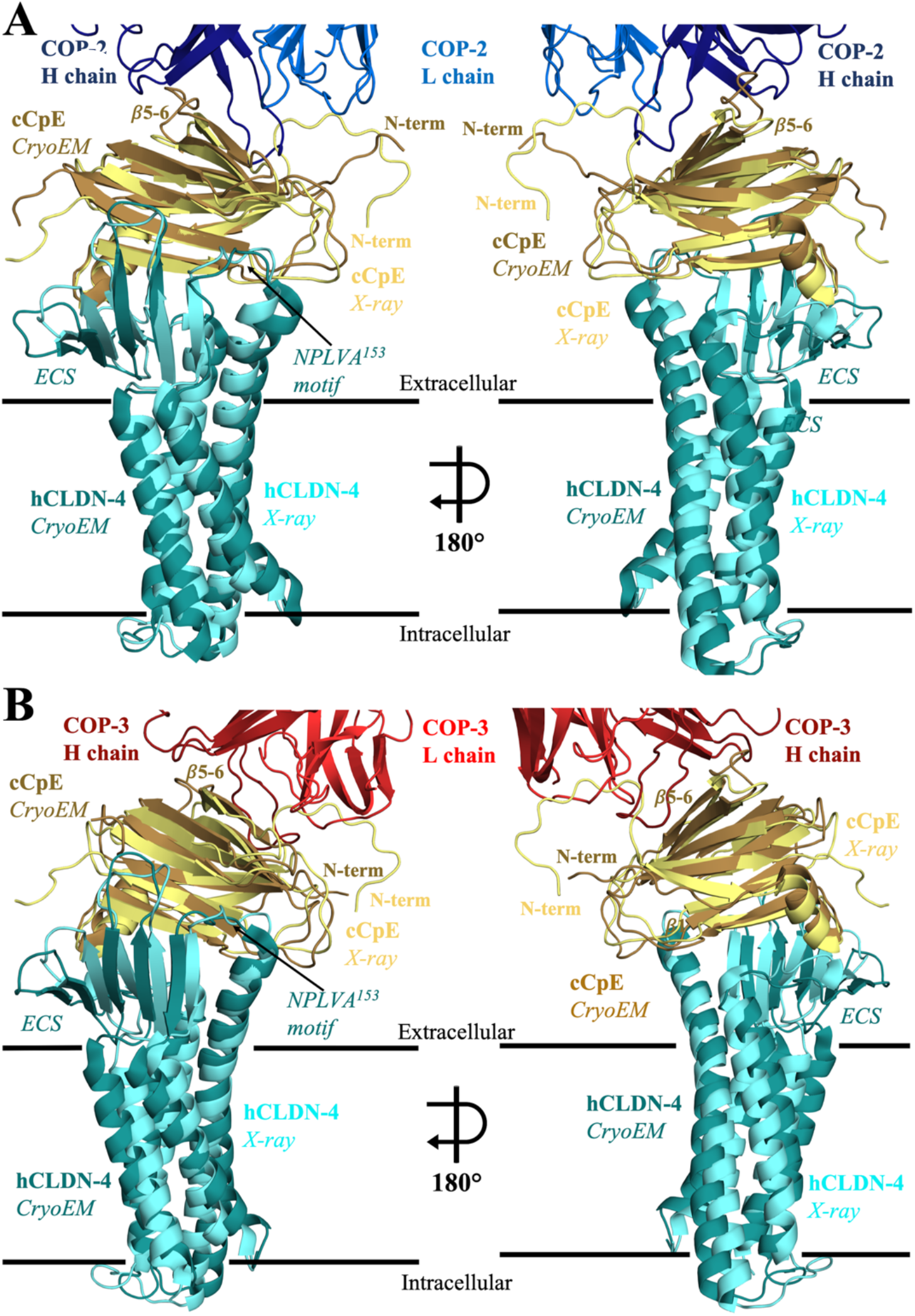
Comparison of Claudin-4/cCpE CryoEM and Crystal Structures. Structural overlays depicting similarity between human claudin-4 (hCLDN-4, teal)/cCpE (sand) complexes from cryoEM, and claudin-4/cCpE (cyan/yellow) complex from X-ray crystal structure PDB ID 7KP4. (**A**) Overlay from COP-2-bound (blue) claudin-4/cCpE complex. (**B**) Overlay from COP-3-bound (red) claudin-4/cCpE complex. Proteins are shown as cartoons with model membrane borders shown as black lines.

**SI Figure S7.**
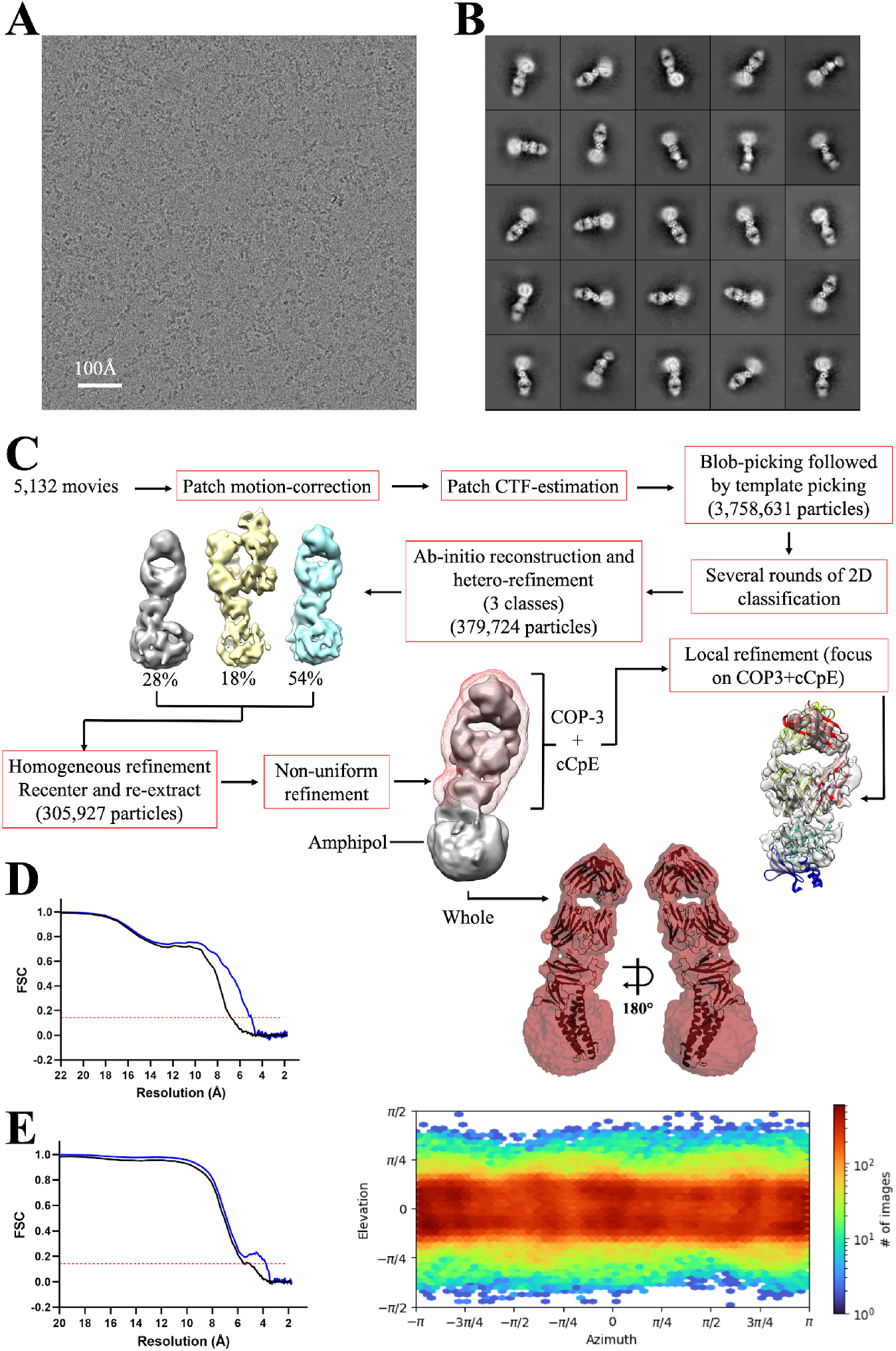
CryoEM Data Processing for Claudin-4/cCpE/COP-3 Complexes. (**A**) Electron micrograph showing the distribution of claudin-4/cCpE/COP-3 complexes in ice. (**B**) Representative 2D class averages of claudin-4/cCpE/COP-3 complexes showing the clear binding of the COP-3 sFab, and signal for TMs within the amphipol belt. (**C**) CryoEM data processing workflow. (**D**) Fourier Shell Correlation (FSC) curves from gold-standard refinement of the entire complex. The 0.143 FSC cutoff is indicated by a dashed red line. The black line indicates FSC without a mask applied, and the blue line indicates FSC with a tight mask applied around the protein and amphipol belt. (**E**) Fourier Shell Correlation (FSC) curves from local refinement of the cCpE/COP-3 region. The 0.143 FSC cutoff is indicated by a dashed red line. The black line indicates FSC without a mask applied, and the blue line indicates FSC with a tight mask applied around the cCpE/COP-3 complex. (**F**) Angular distribution of particles in the final refinement.

**SI Figure S8.**
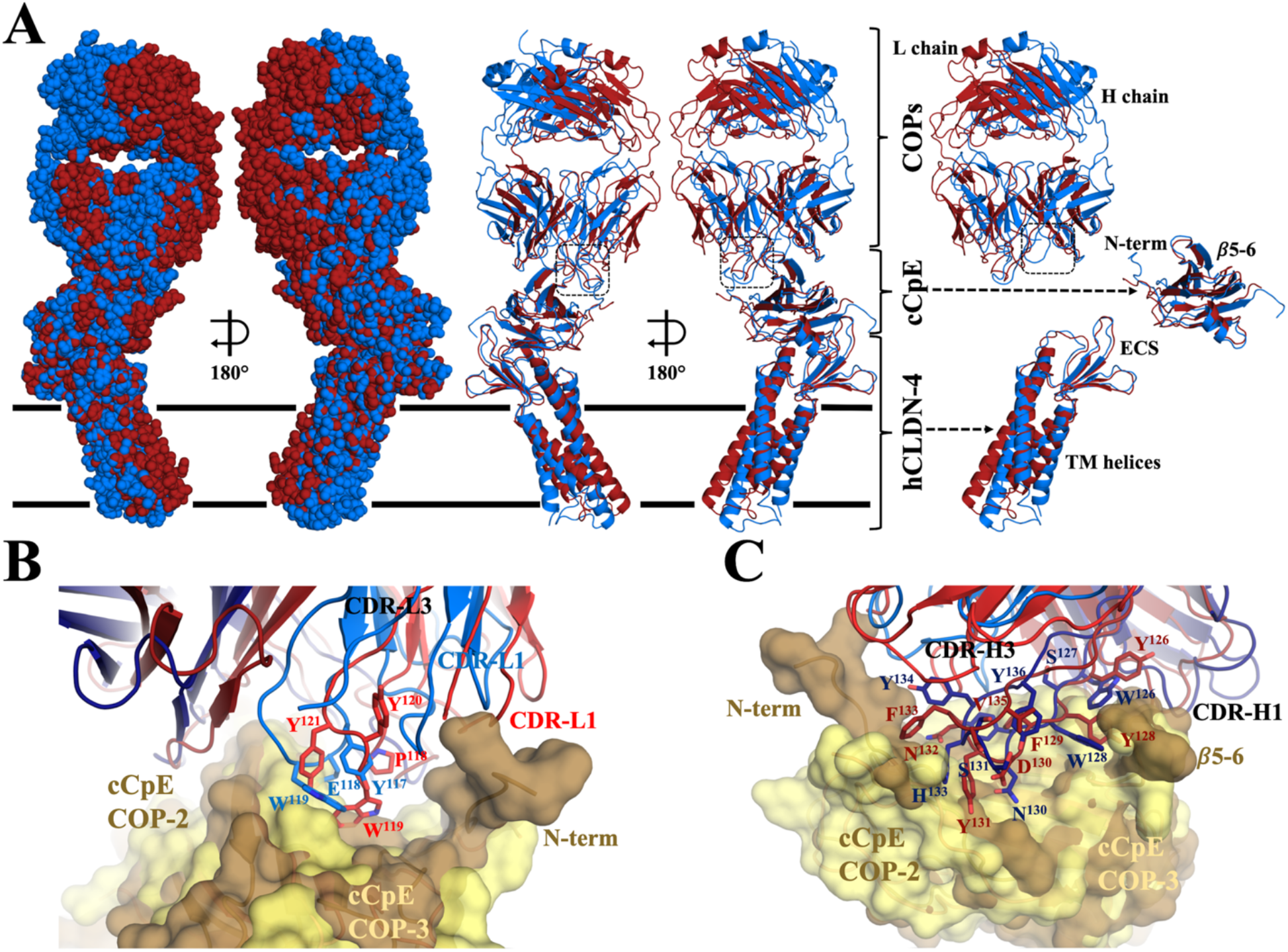
Comparison of Claudin-4/cCpE/COP-2 and Claudin-4/cCpE/COP-3 Structures. Structures were overlaid using Chimera (46). (**A**) Overlay of COP-2-bound (blue) and COP-3-bound (maroon) human claudin-4 (hCLDN-4)/cCpE complexes depicted as a surface (left) or cartoon (middle). Model membrane borders shown as black lines. Overlays of each protein component were made by removing the other two components for ease of visualization and comparison (left). (**B**) Overlay of COP-2 L chain (blue) bound to cCpE (copper) and COP-3 L chain (red) bound to cCpE (yellow). COPs are represented as cartoons while cCpE is shown as a semi-transparent surface. (**C**) Overlay of COP-2 H chain (dark blue) bound to cCpE (copper) and COP-3 H chain (maroon) bound to cCpE (yellow). Proteins are shown as in **B**.

**SI Figure S9.**
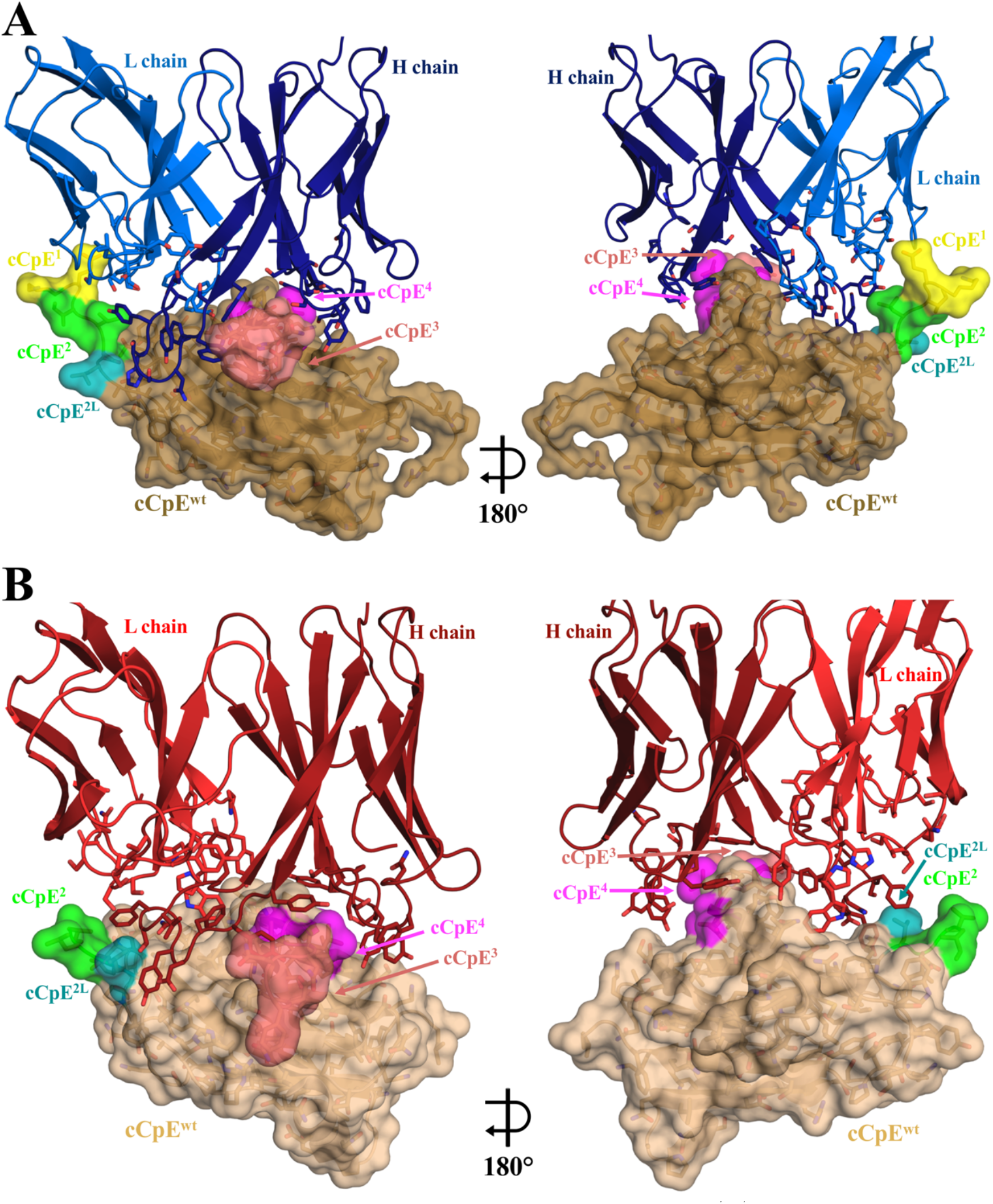
Modelled Changes to COP Surface Epitopes for cCpE^mutants^. (**A**) COP-2 (blue) bound to cCpE^wildtype^ (copper) with areas of cCpE^mutants^ colored as follows: mutant cCpE^1^ (yellow), cCpE^2^ (green), cCpE^2K^ (dark green), cCpE^2L^ (teal), cCpE^3^ (salmon), and cCpE^4^ (magenta). (**B**) COP-3 (red) bound to cCpE^wildtype^ (tan) with areas of cCpE^mutants^ colored as in **A**. The cCpE^mutants^ are visualized as in **SI Fig. S8**. COPs are shown as cartoons while cCpEs are shown as surfaces.

**SI Figure S10.**
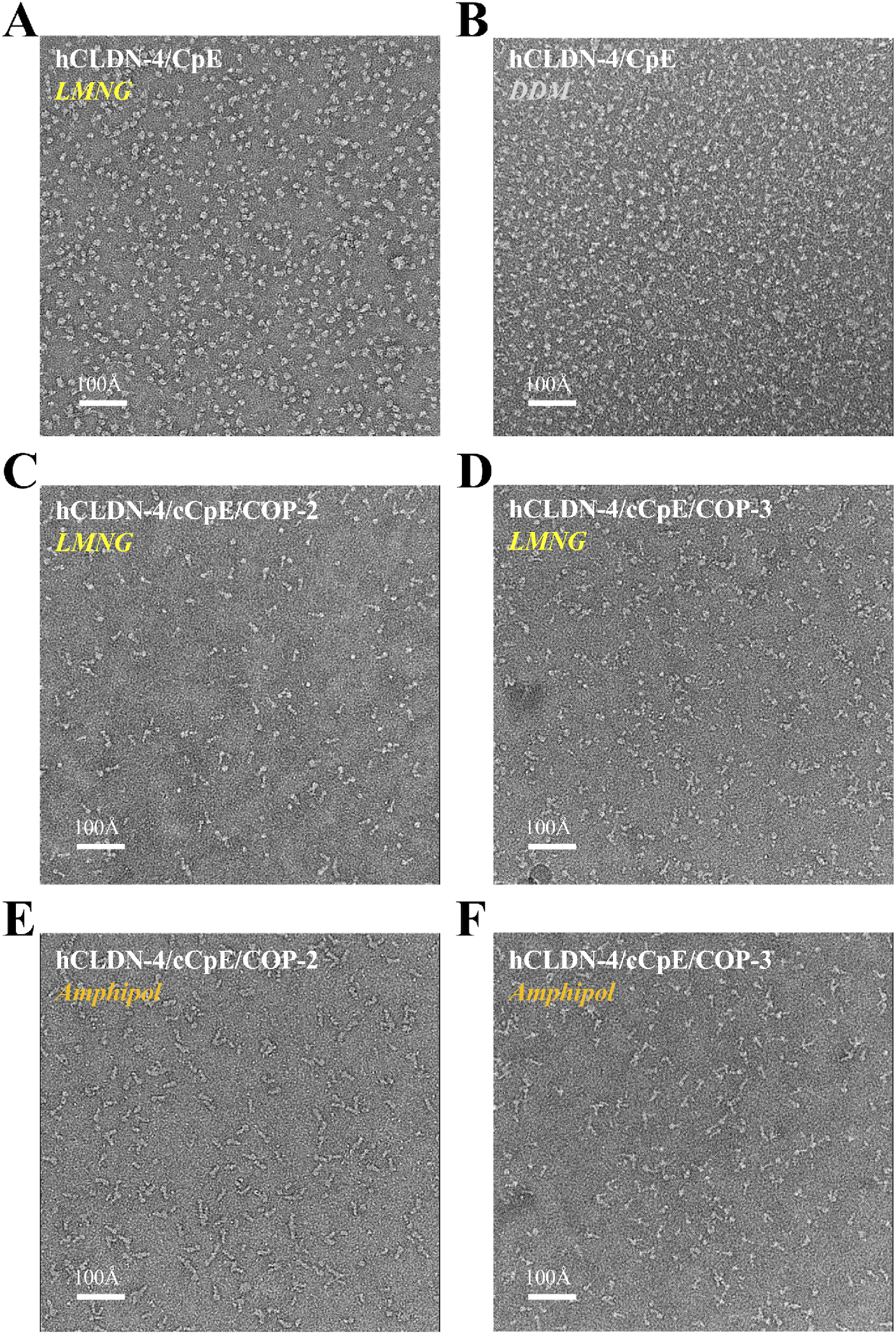
Negative Stain EM of Claudin-4/Enterotoxin Complexes in Various Mimetics. (**A**) Human claudin-4 (hCLDN-4) solubilized in LMNG bound to CpE. (**B**) hCLDN-4solubilized in DDM bound to CpE. (**C**) hCLDN-4 solubilized in LMNG bound to cCpE and COP-2. (**D**) hCLDN-4 solubilized in LMNG bound to cCpE and COP-3. (**E**) hCLDN-4 solubilized in amphipol bound to cCpE and COP-2. (**F**) hCLDN-4 solubilized in amphipol bound to cCpE and COP-3. All negative stain data were collected at 200 keV using a screening cryoEM microscope. Samples from panels **C** and **F** were collected and further processed, resulting in the structures presented here.

## Notes

### Competing Interest Statement

The authors have declared no competing interest.

